# scRNA-seq mixology: towards better benchmarking of single cell RNA-seq analysis methods

**DOI:** 10.1101/433102

**Authors:** Luyi Tian, Xueyi Dong, Saskia Freytag, Kim-Anh Lê Cao, Shian Su, Abolfazl JalalAbadi, Daniela Amann-Zalcenstein, Tom S. Weber, Azadeh Seidi, Jafar S. Jabbari, Shalin H. Naik, Matthew E. Ritchie

**Affiliations:** The Walter and Eliza Hall Institute of Medical Research, 1G Royal Parade, Parkville, VIC 3052, Australia; Department of Medical Biology, The University of Melbourne, Parkville, VIC 3010, Australia; College of Life Science, Zhejiang University, 866 Yuhangtang Road, Hangzhou, Zhejiang Province, 310058, P.R. China; Harry Perkins Institute of Medical Research, Nedlands, WA 6009, Australia; Melbourne Integrative Genomics, School of Mathematics and Statistics, The University of Melbourne, Parkville, VIC 3010, Australia; Australian Genome Research Facility, Level 13, Victorian Comprehensive Cancer Centre, 305 Grattan Street, Melbourne, VIC 3000, Australia

## Abstract

Single cell RNA sequencing (scRNA-seq) technology has undergone rapid development in recent years, bringing with new challenges in data processing and analysis. This has led to an explosion of tailored analysis methods for scRNA-seq data to address various biological questions. However, the current lack of gold-standard benchmark datasets makes it difficult for researchers to systematically evaluate the performance of the many methods available. Here, we designed and carried out a realistic benchmark experiment that included mixtures of single cells or ‘pseudo cells’ created by sampling admixtures of cells or RNA from up to 5 distinct cancer cell lines. Altogether we generated 14 datasets using droplet and plate-based scRNA-seq protocols, compared multiple data analysis methods in combination for tasks ranging from normalization and imputation, to clustering, trajectory analysis and data integration. Evaluation across 3,913 analyses (methods × benchmark dataset combinations) revealed pipelines suited to different types of data for different tasks. Our dataset and analysis present a comprehensive comparison framework for benchmarking most common scRNA-seq analysis tasks.

---

The rapid development of transcriptomic technology for single cell analysis has created a need for systematic benchmarking in order to understand the strengths and weaknesses of different computational methods. To date, there have been several comparison studies of different protocols and computational methods for single cell RNA sequencing (scRNA-seq). Tung *et al*. [47] assessed the batch variation in scRNA-seq data and highlighted the importance of experimental designs that avoid confounding of biological and technical effects. The performance of particular scRNA-seq data analysis methods have been evaluated for tasks including normalization [6], feature selection [51], differential gene expression analysis [43], clustering [8, 9] and trajectory analysis [39]. These studies compare methods using either experimental data where cell type labels are available or simulated datasets. Such ground truth is however imperfect for reasons outlined below, and simulations rely on assumptions that may not reflect the true nature of scRNA-seq data. Also, by focusing only on specific tasks, these studies lack a complete picture of performance at the pipeline level of scRNA-seq data analysis.

Considering the heterogeneity between scRNA-seq datasets in terms of the number of clusters (cell types/states) and the presence of various technical artifacts, we set out to design a realistic gold-standard scRNA-seq control experiment that combines ground truth with varying levels of biological complexity. Two strategies are commonly employed to create such gold-standard gene expression datasets. The first uses small collections of exogenous spike-in controls (such as ERCCs [21]) that vary in expression in a predictable way and have been widely adopted in scRNA-seq studies [45]. The second involves either the dilution of RNA from a reference sample or mixing of RNA or cells from two or more samples to induce systematic genome-wide changes. An early example of an scRNA-seq control dataset was presented in Brennecke *et al*. [3] and involved a dilution series to explore sensitivity of the Smart-seq protocol. Grün *et al*. [11] generated a benchmark dataset using single mouse embryonic stem cells (mESC) together with bulk RNA extracted from the same population, diluted to single cell equivalent amounts to quantify biological and technical variability. A limitation of these experiments is their lack of biological heterogeneity which makes them less useful for comparing analysis methods. Mixture designs, in which RNA or cells are mixed in different proportions to generate biological heterogeneity with in-built truth have been successfully used to benchmark microarray [7], RNA-seq [42].

To combine the strengths of these approaches, we designed a series of experiments using mixtures of either cells or RNA from up to 5 cancer cell lines and included a dilution series to simulate variations in the RNA content of different cells as well as ERCC spike-in controls wherever possible. Data were generated across four single-cell platforms (CEL-seq2, SORT-seq, 10X Chromium and Drop-seq). Our scRNA-seq mixology design simulates varying levels of biological noise, with sample sizes varying from around 200 cells to 4,000 cells and known population structure to allow benchmarking of different analysis tools.

In this article we specifically highlight data normalization and imputation, clustering, trajectory analysis and data integration to showcase the broad range of tasks that our unique collection of datasets allows us to benchmark. We chose popular methods for each task that use different algorithms and are mostly implemented in R for convenience. Using a novel software platform *(CellBench*), we examine 3,913 analyses representing different combinations of methods across these datasets to assess performance of various pipelines. Our analyses across multiple datasets allows evaluation of the generalizability of different methods and their combinations to help inform best practice in scRNA-seq data analysis.

## Results

### scRNA-seq mixology provides ground truth for benchmarking

As summarised in Supplementary Table 1, the scRNA-seq benchmarking experiment spanned 2 plate-based and 2 droplet-based protocols and involved 3 different experimental designs with replicates, yielding 14 datasets in total. Our experiment used the 5 human lung adenocarcinoma cell lines H2228, H1975, HCC827, A549 and H838. The experimental design involved either mixtures of RNA or single cells from these cell lines and single cells. For the *single cell* designs, 3 cell lines (H2228, H1975, HCC827) were mixed equally and processed by 10X Chromium, Drop-seq [30] and CEL-seq2 [14] (referred to as *sc*_*10X, sc*_*Drop-seq* and *sc*_*CEL-seq2*, respectively). Similarly, 5 cell lines were mixed equally and processed by 10X Chromium and CEL-seq2 (referred to as *sc*_*10x*_*5cl* and *sc*_*CEL-seq2*_*5cl*_*p1, sc*_*CEL-seq2*_*5cl*_*p2* and *sc*_*CEL-seq2*_*5cl*_*p3* for three plates respectively). For the *‘pseudo cell’* designs, we used plate-based protocols to mix and dilute samples in 2 different ways. First, we created 9-cell mixtures by sorting different combinations of cells from 3 cell lines (H2228, H1975, HCC827) and generating libraries using CEL-seq2. The material after pooling from 384 wells were sub-sampled to either 1/9 or 1/3 of the total mixture to simulate cells with varying mRNA content and using different PCR product clean up ratios (sample:beads) ranging from 0.7:1 to 0.9:1. These data are referred to as *cellmix1* to *cellmix4* (Supplementary Figure 1B; Supplementary Table 1). For each mixture, we also created a reference comprised of mixtures of 90 cells for each mixture (referred to as *cellmix5*). The second design created *‘pseudo cells’* by mixing bulk RNA obtained from 3 cell lines (H2228, H1975, HCC827), which were diluted to create single cell equivalents (varying from 3.75, 7.5, 15 to 30 pg per well) to again create controlled variations in RNA content. Data were generated for this *RNA mixture* design using CEL-seq2 and SORT-seq [32] (referred to as *RNAmix*_*CEL-seq2* and *RNAmix_Sort-seq*, Supplementary Figure 1A; Supplementary Table 1).

**Figure 1.**
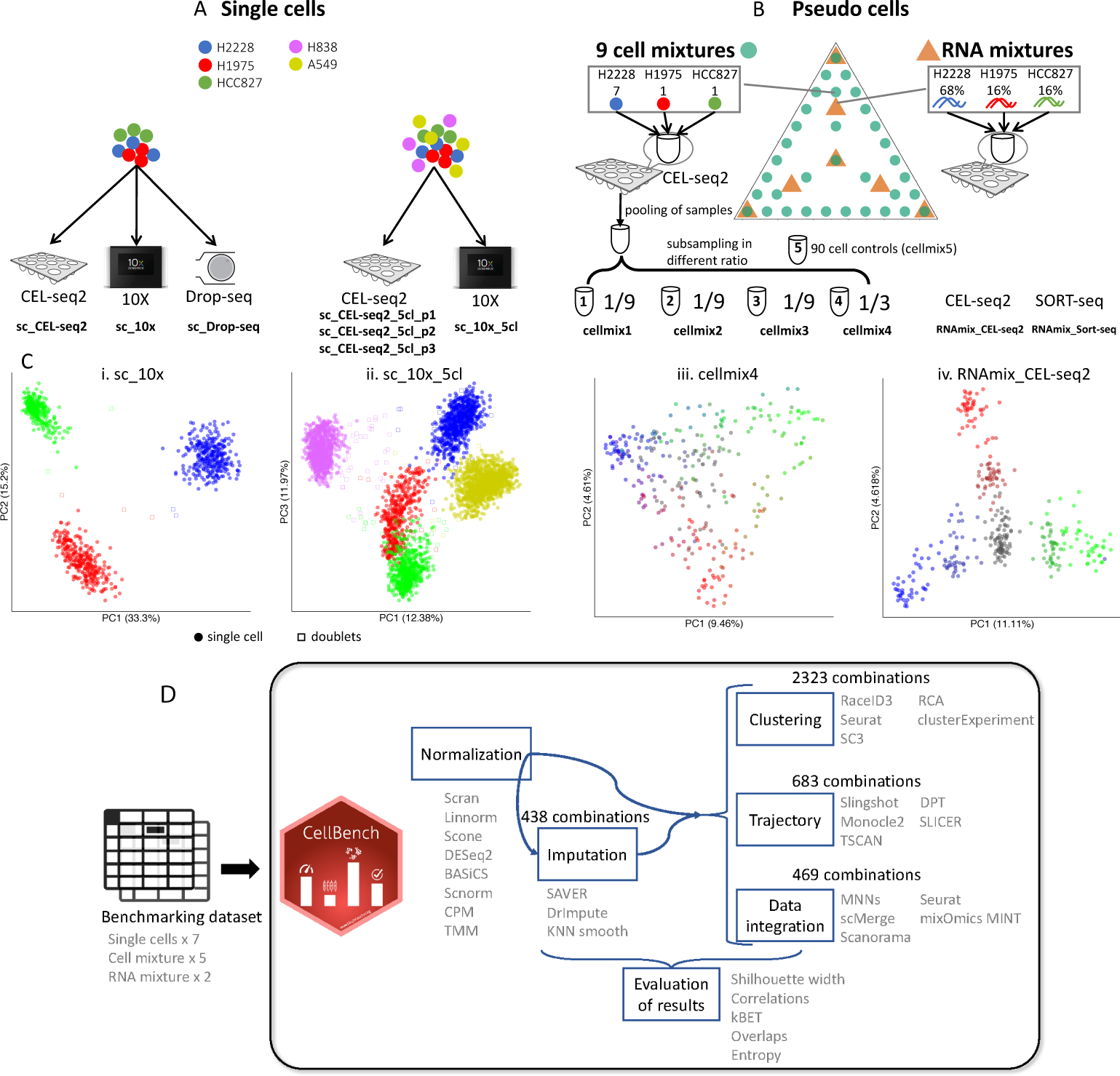
Overview of our scRNA-seq mixology experimental design. The benchmark exper-imental design involving single cells and *‘pseudo cells’*. (**A**) Single cells from three cell lines (H2228, H1975, HCC827) were combined in equal proportions and scRNA-seq was performed using the CEL-seq2 (*sc*_*CEL-seq2*), 10X Chromium protocols (*sc*_*10x*) and Drop-seq (*sc*_*Drop-seq*) protocols. Similarly, equal mixtures of 5 cell lines were processed by CEL-seq2 (*sc*_*CEL-seq2*_*5cl*_*p1, sc*_*CEL-seq2*_*5cl*_*p2* and *sc*_*CEL-seq2*_*5cl*_*p3*) and 10X Chromium (*sc*_*10x*_*5cl*). (**B**) *‘Pseudo cells’* were created by mixing cells or RNAs. For cell mixtures, we sort different combinations of 9-cells from the three cell lines into 384-well plates and subsequently diluting them to obtain single cell equivalent amounts of RNA (*cellmix1* to *cellmix4*). 90-cell mixtures were also include to create pseudo bulk references for each mixture (*cellmix5*). The RNA mixtures were created by mixing RNA that obtained from bulk samples from the three cell lines in different proportions (*RNAmix*_*CEL-seq2, RNAmix_Sort-seq*). (**C**) PCA plots from representative datasets for each design (normalized using *scran*) highlight the structure present in each experiment. The amount of variation explained by each PC is included in the respective axis labels. (**D**) Workflow for benchmarking different analysis tasks. After quality control, each dataset was processed by different normalization and imputation methods. Each processed gene expression matrix was then used as input to various downstream tasks. This resulted in 3,913 different analysis (methods × dataset) combinations that allowed performance assessment at the pipeline level. The *CellBench* R package facilitated these combinatorial analyses.

The three designs incorporate ground truth in various ways. For the *single cell* mixture datasets, the ground truth is the cell line identity which can be determined for each cell based on known genetic variation of each cell line. The *single cell* mixtures were generated using three different technologies, which allows for comparisons of data integration methods. The *cell mixture* and *RNA mixture* datasets contain 34 and 7 groups respectively that give a continuous structure. For the mixture data, the composition of cells/RNA that make up each *‘pseudo cell’* are known, which serves as ground truth. Moreover, the *RNA mixture* dataset contains technical replication and a dilution series, which is ideal for benchmarking normalization and imputation methods that are intended to deal with such technical variability. The data characteristics and analysis tasks each experimental design is best suited to benchmark are summarised in Supplementary Table 2.

By comparing a range of quality control metrics collected across datasets using *scPipe* [46], we observed that the data from all platforms were of consistently high quality in terms of their exon mapping rates and the total unique molecular identifier (UMI) counts per cell (Supplementary Figure 2). Interestingly, we found substantial differences in the percentage of reads mapping to intron regions in datasets generated from different protocols and experimental designs (Supplementary Figure 2A). After normalization by *scran*, the Principal Component Analysis (PCA) plots from four representative datasets show that our single cell and mixture datasets successfully recapitulate the expected population structure induced by our design (Figure 1C; t-SNE and UMAP visualisations provided in Supplementary Figure 3A).

**Figure 3.**
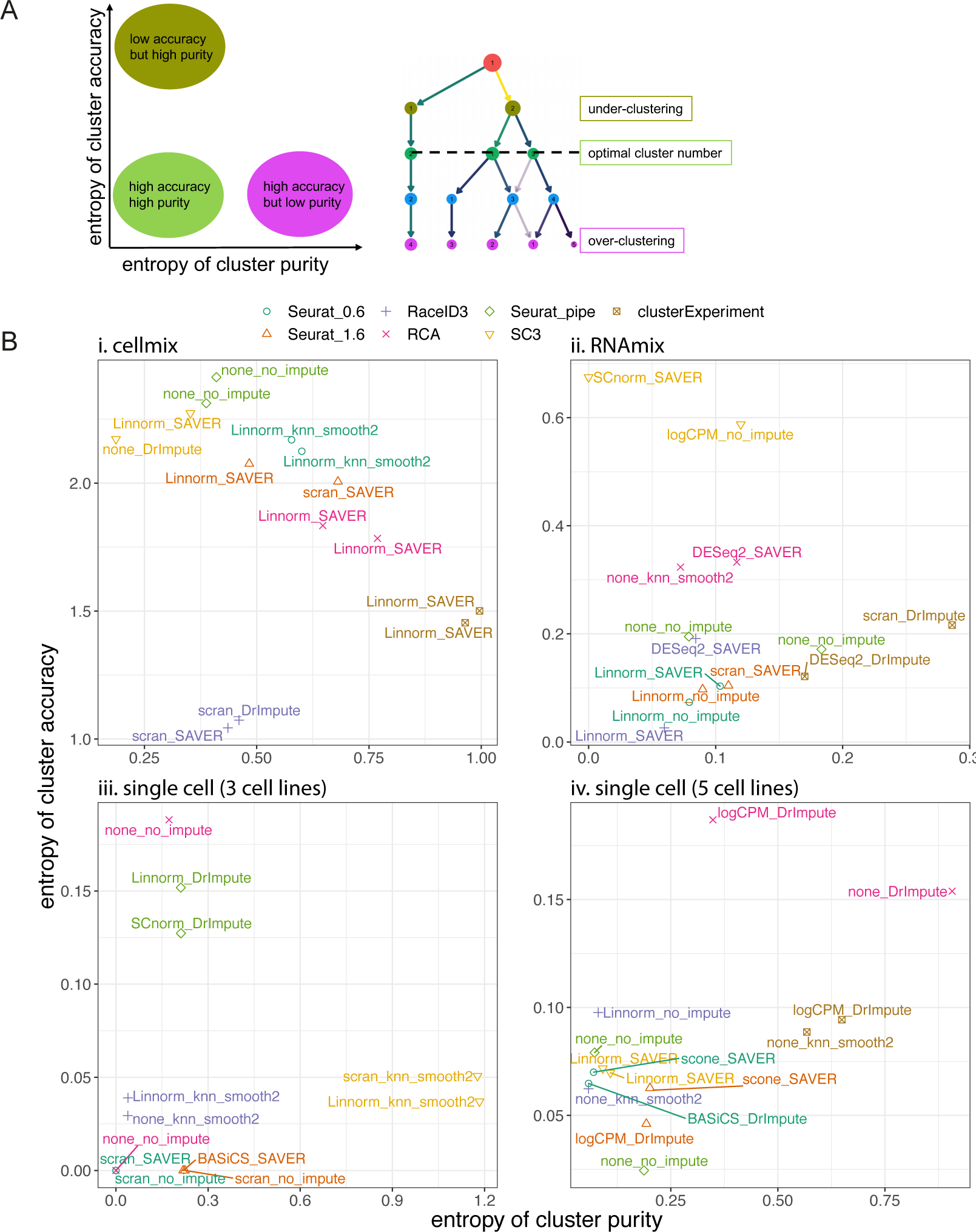
Comparisons of scRNA-seq clustering methods. (**A**) An overview of the evaluation approach. Entropy of cluster accuracy measures the degree of over-clustering, while Entropy of cluster purity measures under-clustering. The clustering tree adapted from the package *Clustree* [53] allows conceptual visualisation of the two measurements. (**B**) Entropy of cluster purity and entropy of cluster accuracy for the top performing results for each method. Colours denote different clustering methods and labels indicate the combination of normalization and imputation methods used as input to the clustering algorithms).

**Figure 2.**
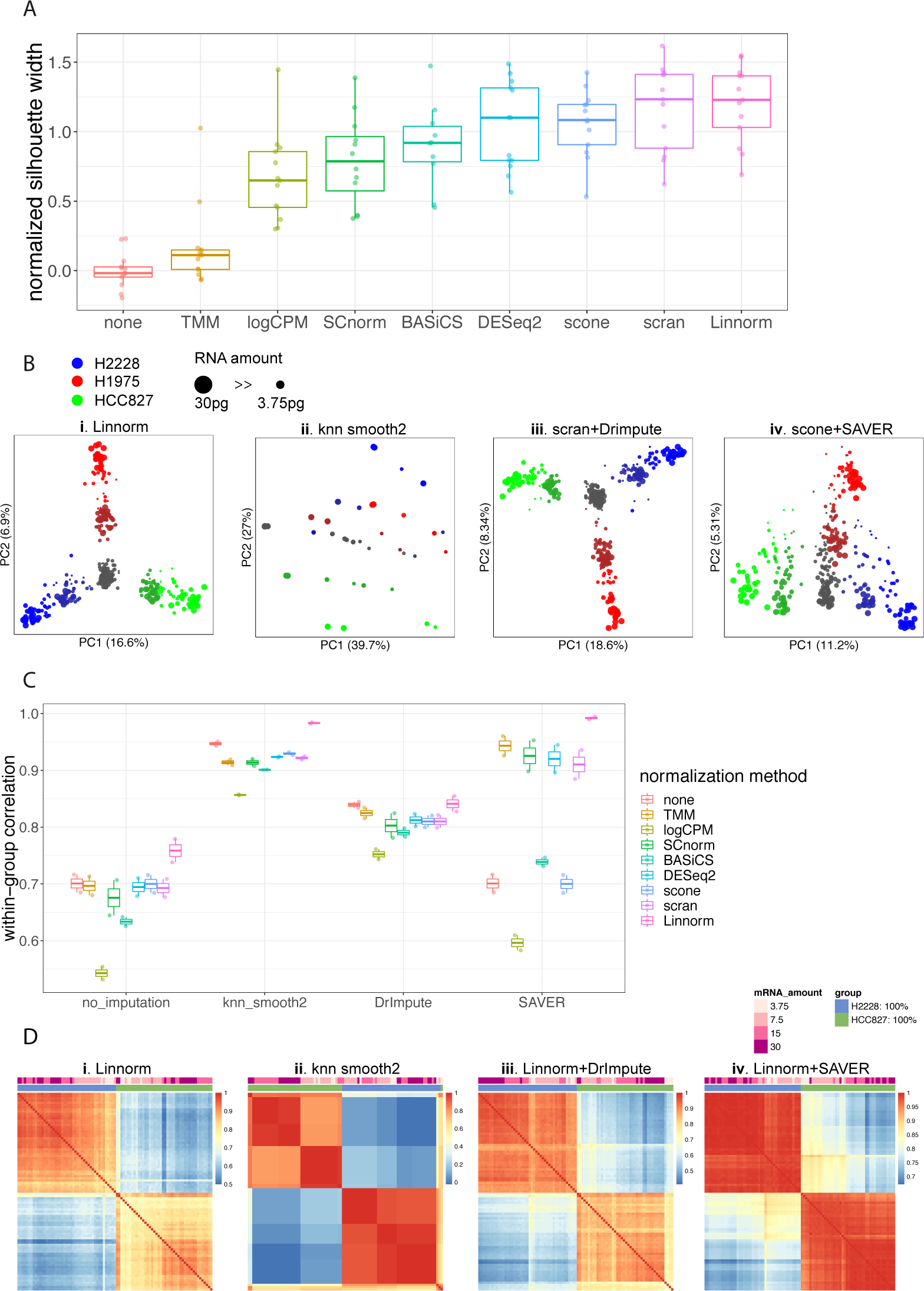
Comparisons of normalization and imputation methods using multiple mixture datasets. (**A**) Silhouette widths calculated using the known cell/mixture groups after different normal-ization methods, summarised across all datasets and normalized against the Silhouette widths obtained without normalization (‘none’). (**B**) Example PCA plots after normalization or with imputation by different methods using the *RNAmix*_*CEL-seq2* dataset. The amount of variation explained by each PC is included in the respective axis labels. (**C**) Average Pearson correlation coefficients for *‘pseudo cells’* within the same groups in the *RNAmix*_*CEL-seq2* dataset for different combinations of normalization and imputation methods. (**D**) Heatmap of Pearson correlation coefficients of samples that have pure H2228 or HCC827 RNA obtained from different imputation methods, clustered by *hclust*.

A summary of the method comparisons performed using these data and the *CellBench* R package is shown in Figure 1D. After quality control, each dataset was processed using different combinations of normalization and imputation methods. Next, each normalized and imputed gene expression matrix was used as the input for several downstream steps, including clustering, trajectory analysis and data integration. The performance of each method was evaluated using different metrics tailored to the ground truth available.

### Comparisons of normalization and imputation methods

Normalization is an important step in the analysis of scRNA-seq data, with the general goal of removing technical noise while retaining biological signal. Imputation on the other hand recovers missing data due to dropout events, which are excess zero counts caused by the limited capture efficiency of scRNA-seq protocols. We evaluated 8 popular normalization methods, including methods developed primarily for bulk RNA-seq such as *TMM* [36], *CPM* [35] *and DESeq2* [28], and others tailored for scRNA-seq, including *scone* [6], *BASiCS* [48], *SCnorm* [2], *Linnorm* [52] and *scran* [29]. Three imputation methods, including *kNN-smoothing* [49], *DrImpute* [10] *and SAVER* [18] were also evaluated using input data normalized by different methods. Benchmarking was performed across 438 analyses representing combinations of normalization and imputation methods.

Performance was evaluated using 2 metrics: the silhouette width of clusters for all datasets, and the Pearson correlation coefficient of normalized gene expression within each group for the *RNA mixture* data, where the group is defined by the known cell line mixing proportions. The silhouette width was calculated based on the PCA results obtained for each method with the cluster identities defined according to the experimental design. Although a wide range of silhouette widths was observed amongst different methods in different datasets, most of the normalization methods, apart from *TMM*, significantly increased silhouette width compared to the unnormalized data (Figure 2A, Supplementary Figures 4A and 4B). Methods tailored to single cell data tend to perform better than methods designed for bulk RNA-seq analysis, with the exception of *DESeq2*, which generated good results. Across all the methods compared, *Linnorm* was the top performer on average, followed by *scran* and *scone*.

**Figure 4.**
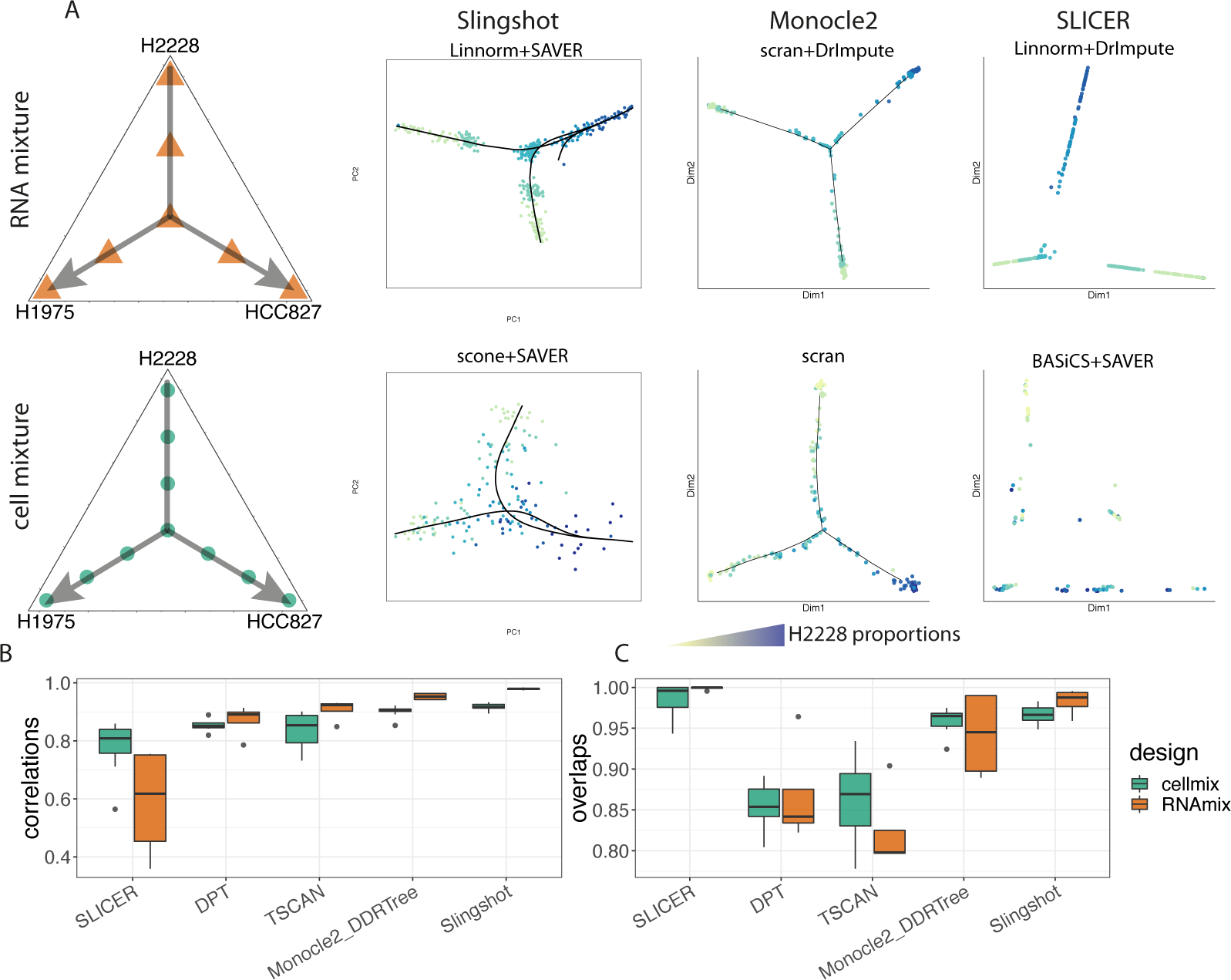
Comparisons of scRNA-seq trajectory analysis methods. (**A**) The trajectory path chosen for the *RNA mixture* dataset (top) and cell mixture dataset (bottom) along with visualisations of the output from *Slingshot, Monocle-DDRTree* and *SLICER*. Cells are coloured by the proportion of H2228 RNA present, which was chosen as the root of the trajectory. (**B**) Boxplot showing the Pearson correlation coefficient between the calculated pseudotime and the ground-truth, for the best performing combination of each method on each dataset. (**C**) The proportion of cells that are correctly assigned to the trajectory.

The *RNA mixture* experiment included pseudo cells with varying amounts of input RNA to simulate dropout events, making it an ideal dataset for evaluating the performance of imputation methods. In general, imputation induces higher intra-group correlation, although considerable differences are observed depending on the normalization method chosen (Figure 2C). *KNN-smoothing* and *DrImpute* show similar results with different normalization methods. *SAVER* has greatest variation in performance, with either the best or worst depending upon the input normalization method applied (Figure 2B-iv and 2C).

In addition to examining the ability of different combinations of methods to recover true signal from noisy data, we also investigated whether spurious cell states could be introduced during imputation. We chose cells from two distinct groups (pure H2228 and HCC827) in the *RNA mixture* dataset and looked at the correlations among samples. Overall, the sample correlation within the same group is lower when the mRNA amount decreases (Figure 2D-i), and is recovered by imputation (Figure 2D-ii-iv). All three methods clearly separate the two pure groups, however *kNN-smoothing* introduces a spurious intra-group correlation structure that is also shown in the PCA of the imputed data (Figure 2B-ii), which is similar to what has been found in another recent study [1]. Moreover, we found the extra clusters were related to the input RNA amount, which implies that *kNN-smoothing* is sensitive to dropout events.

### Comparisons of clustering methods

The benchmark datasets we designed varied in the expected cluster number (3 or 5 for the single cell dataset, 7 for the*RNA mixtures* and 34 for the *cell mixtures*) and the degree of cluster separability (low for the *cell mixtures*, moderate for the *RNA mixtures* and high for the *single cell datasets*. This allowed us to assess clustering performance in a variety of settings. Five representative methods, including *RaceID3* [15], *RCA* [25], *Seurat* [30], *clusterExperiment* [33] and *SC3* [23], were evaluated across all datasets. As there is no function to choose the optimal number of clusters in *Seurat*, two resolutions, 0.6 and 1.6 were used. The resolution parameter controls the number of clusters, with higher values producing more clusters. In addition to the various normalized gene expression matrices, we also evaluated *Seurat* using its own default pipeline that starts from the raw gene count matrix (denoted *Seurat pipe*).

We measured the performance of clustering methods by calculating both the entropy of cluster accuracy (ECA) and entropy of cluster purity (ECP). After clustering, we have both the cluster labels assigned by the different methods, and the known group labels that provide us with ground truth. The entropy of cluster accuracy measures the diversity of the true group label within each cluster assigned by the clustering algorithm. ECA does not account for over-clustering, and in an extreme case, a method that assigns a unique cluster for each cell will have an ECA of 0. In contrast, the entropy of cluster purity measures the diversity of the calculated cluster labels within each of the true groups and offers no control of under-clustering (a method that assigns all cells into one cluster will have an ECP of 0, which is indicative of high cluster purity). Therefore, we consider these two metrics together to account for both under and over-clustering, with methods that have both low ECP and low ECA having optimal cluster assignments (Figure 3A). We found good correlation between these two metrics and the Adjusted Rand Index (ARI) [19], which is a commonly used metric to evaluate clustering performance by computing the similarity to the annotated clusters (Supplementary Figure 5A).

**Figure 5.**
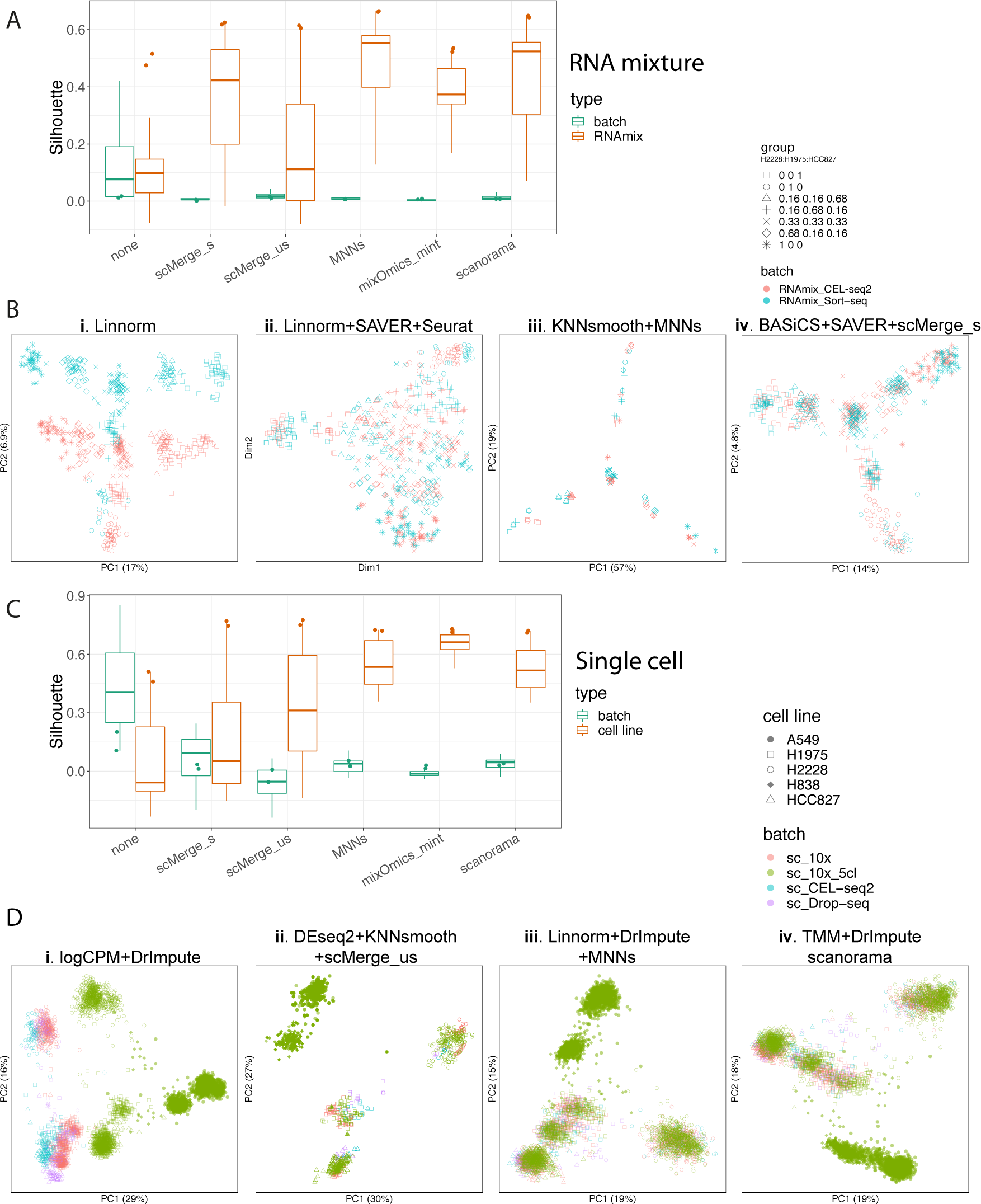
Comparisons of data integration methods for batch effect correction for the *RNA mixture* (A, B) and the 4 *single cell* experiments (C, D). (**A**,**C**) Silhouette width calculated on either batch information or known sample group. Input data were based on a combination of different normalization and imputation methods, with top performing combination indicated for each method. (**B**,**D**) Examples of dimension reduction visualisations for some data integration methods (ii, iii, iv). *Seurat* results were visualised with t-SNE and other methods’ results with PCA. *scMerge*_*s*: supervised *scMerge*; *scMerge*_*us*: unsupervised *scMerge*.

As shown in Figure 3B, which presents the best two results for each method, no particular algorithm consistently outperformed others across all experimental designs under default settings. In general, *Seurat* achieved a good balance between under-clustering and over-clustering across all datasets, and performs best when there was clear separation between cell types, as was the case in the single cell datasets (Figure 3B-iii,iv). *RaceID3* outperformed all other methods in the more complex cell mixture datasets. The accuracy of all methods was lowest in the *cell mixture* datasets (Figure 3D), due to the continuous population structure which gives less separation between different clusters compared to the other datasets. For the single cell datasets, we note that the true cluster number given by genotype information is likely to be an underestimate, with subtle sub-clusters present within each cell line, as highlighted in the t-SNE and UMAP visualisations (Supplementary Figure 3Ai-ii). Methods that over-cluster the single cell data may well capture true biological signal.

In addition to showing the best results for each method, a linear model was fitted to 2,323 analysis combinations (different normalization, imputation and clustering methods applied across different datasets) using either ARI or the ‘true’ number of clusters as dependent variables and the different methods as covariates (Supplementary Figures 5A and 5B) to further investigate the contribution of each method to the results. Similar to what has been shown in Figure 3B, the *clusterExperiment* method frequently failed to recover the expected population structure, and *SC3* under-clusters most datasets. Interestingly, we found that *kNN-smoothing* was associated with lower ARI and higher cluster numbers, which is consistent with our previous results (Figure 2D-ii), and serves as a further indication that this method can readily introduce spurious clusters.

### Comparisons of trajectory analysis methods

Five methods, including *Slingshot* [44], *Monocle2* [34], *SLICER* [50], *TSCAN* [20] *and DPT* [12] *were evaluated using the RNA mixture* and *cell mixture* datasets. These datasets were chosen as they both contain clear *‘pseudo trajectory’* paths from one pure cell line to another that are driven by controlled variations in RNA amount. For simplicity, we chose H2228 as the root state of the trajectory (Figure 4A). We evaluated the correlation between the pseudotime generated from each method and the rank order of the path from H2228 to the other cell lines based on the RNA mixture information (Figure 4B) to examine whether each method can position cells in the correct order. In addition, we calculated the coverage of the trajectory path (Figure 4C), which is the percentage of cells that have been assigned to the correct path, and assesses the sensitivity of the method. We used data generated from combinations of normalization and imputation methods as input to each trajectory analysis method, to assess their impact on performance. In total, 683 analysis combinations (different normalization, imputation and trajectory analysis methods applied to different datasets) were evaluated.

For each method in each dataset, we selected their best results from all combination based on the performance metrics (Figure 4B, 4C). In addition to that, a linear model was used to characterize the average performance for each method (Supplementary Figure 8A, 8B). *Slingshot* and *Monocle2* showed robust results according to both metrics and generated meaningful representations of the trajectory, while *Slingshot* sometimes gave an extra trajectory path (Figure 4A, Supplementary Figure 6). In contrast, *SLICER* places all cells in the correct path but was unable to order them correctly or recover the expected structure induced by the mixture designs.

**Figure 6.**
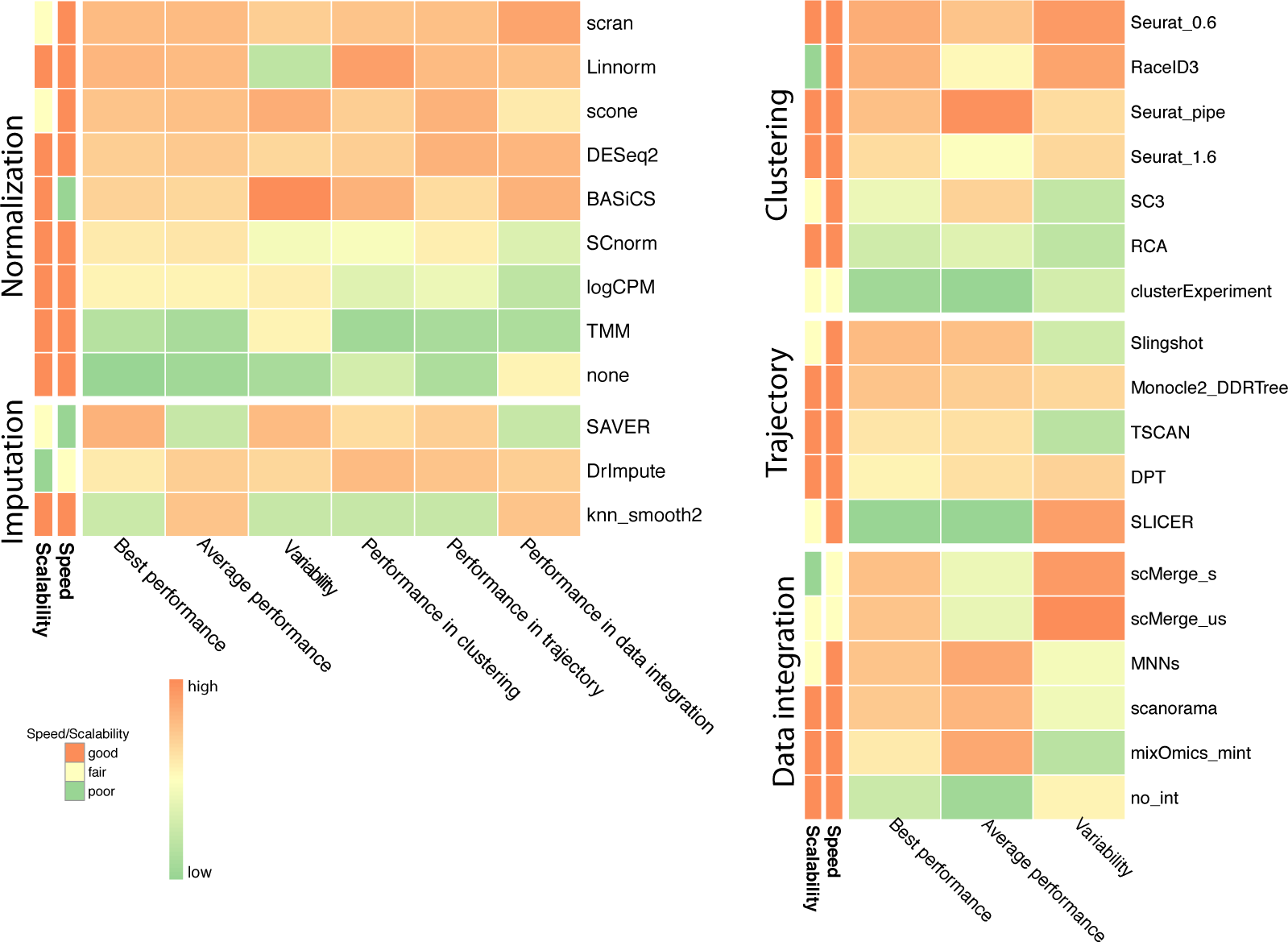
Summary of results from methods comparisons using scRNA-mixology datasets. Methods are ranked by the best performance in each category. The average performance and variability show the average result and the variance when given different input dataset and processed by different upstream methods. For normalization and imputation, the impact each method have on downstream analysis is also shown, which measures the changes in results when applying such method. Results have been scaled and standardised to apply to the same colour scale.

Despite the similar performance of *Slingshot* and *Monocle2*, their results differ in terms of the way they position the cells. *Slingshot* does not perform dimensionality reduction itself and presents the result as is, whereas *Monocle2* uses DDR-tree for dimensionality reduction, and tends to place cells at the nodes of the tree rather than in transition between two nodes (Figure 4A). For example, the *RNA mixture* dataset has 7 clusters by design which are equally distributed along the path between one pure cell line and another. *Monocle2* assigns most of the cells to the three terminal states, leaving only a few in between, which does not reflect the designed structure. Indeed, this feature might fit real situations in cell differentiation, where most cells are in defined cell states with only a small proportion in transition between different groups. However, such an assumption may not always hold and care is therefore needed when interpreting the results.

### Comparisons of data integration methods

Whilst combining scRNA-seq data between studies is an attractive way to increase cell number and ensure reproducible results, there are many challenges to address including high drop-out rates, technical noise introduced during library preparation and variations in sequencing depth per cell. None of the methods proposed so far have been compared on well-designed benchmark datasets generated using multiple protocols. We used the *single cell* (*sc*_*CELseq2, sc*_*10X, sc*_*Dropseq* and *sc*_*10x*_*5cl*) and *RNA mixture* (*RNAmix*_*CELseq2* and *RNAmix_Sortseq*) experiments to compare state-of-art methods including *MNNs* [13], *Scanorama* [16], *scMerge* [27], *Seurat* [4] *and MINT* [37]. The 5 cell line CEL-seq2 datasets (*sc*_*CELseq2*_*5cl*_*p1, sc*_*CELseq2*_*5cl*_*p2* and *sc*_*CELseq2*_*5cl*_*p3*) were excluded due to their high doublet rates (Supplementary Figure 3B). As expected, when naively combining the independent datasets (Figure 5B-i, 5D-i), clear separations related to the different protocols were observed in the PCA plots. The input data for each data integration method resulted from different combinations of normalization and imputation methods. Results from 469 different analyses were assessed in total, made up of different normalization, imputation and data integration methods applied across two datasets.

*MNNs, Scanorama* and *scMerge* generate batch-corrected data which can then be analyzed using other downstream analysis tools, while Diagonal Canonical Correlation Analysis combined with Dynamic Time Warping from *Seurat* and *MINT* [37] output a low-dimensional representation of the data. *MINT* includes an embedded gene selection procedure whilst projecting the data into a lower dimensional space. We assessed the methods’ performance with the silhouette width according to protocol and known cell line or mixture group information (Figure 5A, 5C). As *Seurat* uses t-SNE for visualisation and dimension reduction where distances are not preserved, we also used *kBET* [5] to quantify the remaining batch effect variation (Supplementary Figure 7A and 7B). When considering the best normalization and imputation methods as input data, most methods performed similarly for the *single cell* design, with the exception of *Seurat* that had a low *kBET* acceptance rate, Supplementary Figure 7A. We observed differences in performance between methods for the *RNA mixture* design that includes a larger number of groups (7) and a smaller number of cells per group than the *single cell* design. *MNNs* led to the best performance according to silhouette width distance, while *Seurat* had the highest *kBET* acceptance rate in this analysis (Supplementary Figure 7B), suggesting a homogeneous mix of batches consistent with the t-SNE visualisation.

## Discussion

We designed and generated a comprehensive scRNA-seq benchmarking dataset with varying levels of biological noise and in-built ground truth via population structure that ranges from simple to complex. These datasets incorporate various mixture designs processed using multiple scRNA-seq technologies to facilitate comparisons of many different analysis tasks. Our analysis highlights systematic differences in intron reads between protocols, which has not been reported before. To demonstrate the broad utility of these data, we performed systematic methods comparisons for 4 key analysis tasks: normalization and imputation, clustering, trajectory analysis and data integration. To manage the large number of comparisons performed, we developed the *CellBench* R package which allows different methods to be combined into pipelines and evaluated more conveniently. This represents an advance over previous studies that typically focus on a specific analysis task. By incorporating many different combinations of normalization and imputation in downstream analyses, we were able to assess the robustness and variability of the final outputs in light of their inputs.

As summarised in Figure 6, the performance of methods varied across different datasets, with no clear winners in all situations. However, consistently satisfactory results were observed for *scran, Linnorm, DrImpute* and *SAVER* for normalization and imputation; *Seurat* for clustering; *Monocle2* and *Slingshot* for trajectory analysis as well as *MNNs* for data integration. Some normalization and imputation method combinations were also found to give good results in most downstream analysis tasks, such as *Linnorm* and *SAVER*. Variations were also observed in the ability of methods to handle different inputs. Methods, such as *Linnorm* and *SC3* produced relatively consistent results regardless of their input dataset, while others such as *SAVER* was more sensitive to these inputs. By evaluating the results from method combinations across different tasks, we observed a number of interesting trends related to the suitability of different preprocessing methods in downstream analysis. For example, we found that although imputation generally improves the results of clustering and trajectory analyses, it can lead to poor mixing of data from different batches (Supplementary Figure 7D) in data integration analyses.

The average running time for most of the methods compared was under 30 minutes, except for *SAVER* and *BASiCS*. Apart from running time, the scalability for each method could also be evaluated since the sample sizes of our datasets varied from 200 to 4,000 cells. We found some methods, such as *DrImpute* and *RaceID3*, have reasonable running time but poor scalability, which suggests that they may not be suitable for analysing very large scRNA-seq datasets, such as the *Tabula Muris* mouse cell atlas [41].

Interestingly, the various ensemble methods, which combine results from multiple algorithms in a bid to improve performance, did not always outperform individual methods. The ensemble method *SC3* and *clusterExperiment* did not outperform other clustering methods, *scone* also gave mixed results on different datasets. Having multiple benchmark datasets with different numbers of cells, and groups and varying levels of biological noise allowed us to objectively assess performance with different data characteristics.

Our comparison is subject to a number of limitations such as the linear mixture settings, which may not be a realistic model for developmental trajectories where regulatory gene expression may be non-linear and non-systemic. Also methods are mostly compared under default settings, which may not give optimal performance across all datasets.

Our benchmarking study serves as a demonstration of the different types of comparisons that can be performed using these comprehensive designs. The number of methods for each task can be easily expanded using our *CellBench* software for a more in depth analysis of specific tasks. *CellBench* can also be used to explore the effect different choices of starting parameters have on the results. These data can also be used to test the performance of other analysis methods, such as data preprocessing (alignment, UMI deduplication, gene-level quantification), feature selection and differential expression analysis. Our benchmarking platform will benefit future package developers as it allows new methods to be evaluated on the same standards, avoiding ambiguity caused by cherry-picking evaluation datasets. We hope that this study will reinvigorate interest in the important area of benchmark data generation and analysis, providing new insights into current best practice and guide the development of better scRNA-seq algorithms in the future to ensure the biological insights derived from single cell technology stand the test of time.

## Methods

### Study design

Five human lung adenocarcinoma cell lines HCC827, H1975, A549, H838 and H2228 were cultured separately and the same batch was processed in three different ways (Figure 1). Firstly, single cells from each cell line were mixed in equal proportions, with libraries generated using three different protocols: CEL-seq2, Drop-seq with Dolomite equipment and 10X Chromium.

Secondly, single cells from three cell lines HCC827, H1975 and H2228 were sorted from the three cell lines into 384-well plates, with an equal number of cells per well in different combinations. For most of the wells, we sorted 9 cells in total, with different combinations of three cell lines distributed in *‘pseudo trajectory’* paths (Supplementary Figure 1B), where the major trajectory is similar to the *RNA mixture* design while the minor trajectory is the combination that only contains cells from two cell lines instead of three, which is similar in design to our previous study [17]. For the major trajectory, we also included the population control for each combination, which includes 90 cells in total (i.e a large sample) instead of 9, while maintaining the cell combinations from the different cell lines. Apart from the trajectory design, we also varied the cell numbers and qualities to study the data characteristics in these configurations. We included 9 replicates with 3 cells in total with one cell from each cell line, to simulate “small cells”. 20 cells with low integrity identified by PI staining. The 9-cell wells were sub-sampled after pooling to get single cell equivalents of RNA, with three replicates in 1/9 and one in 1/3. We applied different clean up ratios to the three replicates after library generation to induce batch effects of a purely technical nature and study how clean up affects the data.

Thirdly, RNA were extracted in bulk from three cell lines (HCC827, H1975 and H2228), mixed in 7 different proportions and diluted to single cell equivalent amounts (Supplementary Figure 1A). In total, there are 8 mixtures in the plate layout with 49 replicates of each mixture. The mix1 and mix2 samples have the same proportions of the three cell lines 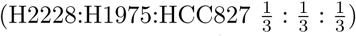 but were prepared separately in order to assess the variation introduced during the RNA dilution and mixture step. In addition to the RNA mixtures, we also designed a dilution series in the same plate to create variations in the amount of RNA added. The amounts ranged from 3.75pg to 30pg (Supplementary Figure 1A-ii) and were intended to simulate differences in cell size. In total, each mixture will have 4 different RNA starting amounts with replicate numbers per mixture of 6:14:14:14 for the 3.75:7.5:15:30 pg group respectively.

### Cell culture and mRNA extraction

The human lung adenocarcinoma cell lines H2228, H1975, HCC827, A549 and H838 were retrieved from ATCC (https://www.atcc.org/) and cultured in Roswell Park Memorial Institute (RPMI) 1640 medium with 10% fetal calf serum (FCS) and 1% Penicillin-Streptomycin. Firstly, three cell lines H2228, H1975 and HCC827 were cultured for the *cell mixture, RNA mixture* and *single cell* experiments. Later, five cell lines (H2228, H1975, HCC827, A549 and H838) were grown separately for another single cell experiment. The cells were grown independently at 37°C with 5% carbon dioxide until near 100% confluency.

For the three cell lines (HCC827, H1975 and H2228), cells were dissociated into single cell suspensions in FACS buffer and sorted for the *cell mixture* and *single cell* experiment (see below for sorting strategy). The remaining cells were centrifuged and frozen at −80°C for later RNA extraction. RNA was extracted using a Qiagen RNA miniprep kit. The amount of RNA was quantified using both Invitrogen Qubit fluorometric quantitation and an Agilent 4200 bioanalyzer to get an accurate estimation. The extracted RNA was then diluted to 60 ng/*µ*l and then mixed in different proportions, according to the study design. The different mixtures were further diluted to create an RNA series that ranged from 3.75pg to 30pg, each of which was dispensed into CEL-seq2 and SORT-seq primer plates using a Nanodrop II dispenser. Prepared RNA mixture plates were sealed and immediately frozen upside down at −80°C.

### Cell sorting and single cell RNA sequencing

For CEL-seq2, single cells were flow sorted into chilled 384-well PCR plates containing 1.2*µ*l of primer/ERCC mix using a BD FACSAria III flow cytometer. Sorted plates were sealed and immediately frozen upside down at −80°C. These plates, together with the RNA mixture plates, were taken from −80°C and processed using an adapted CEL-Seq2 protocol with the following variations. The second strand synthesis was performed using NEBNext Second Strand Synthesis module in a final reaction volume of 8 *µ*l and NucleoMag NGS Clean-up and Size select magnetic beads were used for all DNA purification and size selection steps. For the 9-cell-mixture plates, clean up of the PCR product was performed with 2 × 0.7-0.9 bead/DNA ratio. For the *single cell* and *RNA mixture* plates, two different clean up ratios for the PCR product were used (0.8 followed by 0.9). The choice of clean up ratio was optimized from the QC results of the 9-cell-mixture data and the SORT-seq protocols. For the 5 cell line single cell mixture experiment, the protocol was further optimised by pooling the sample after first strand cDNA synthesis.

The 9-cell-mixture plates were sorted according to the plate design. Each well contained 9 cells in total in different combinations, and was processed using our adapted CEL-seq2 protocol described above with variations in the pooling step. After the second strand synthesis, materials from the 9-cell-mixtures and 90-cell population controls were pooled separately into different tubes and the volumes were measured. Then for the 9-cell-mixture sample, 3 × 1/9 and 1 × 1/3 of the total pooled material were taken and these four samples were processed separately in the following step. At the PCR product clean up stage, the clean up ratios for the 3 × 1/9 samples were 0.7, 0.8 and 0.9 respectively, and 0.7 for the 1/3 9-cell-mixture sample and the 90-cell population controls. The SORT-seq protocol is similar to CEL-seq2 but uses oil to prevent evaporation. This allows reductions in the reaction volume which can be dispensed using the Nanodrop II liquid handling platform (GC biotech). In summary, 2.0*µ*l vapor-lock oil was added to each well of the plate, followed by 0.1*µ*l of primer/ERCC mix. The reaction volume for RT and first strand synthesis are 0.075*µ*l and 0.568*µ*l respectively. The composition of the various mixes was the same as for CEL-seq2. The sample pooling was achieved by centrifuging the plates upside down into a container covered with parafilm and carefully separating the oil from the other materials. The PCR clean up ratio used for SORT-seq was 0.8 followed by 0.9. We experienced significant sample loss during sample pooling such that only 60% of the total volumes were recovered, which is lower compared to the CEL-seq2 protocol (90%).

For the 10X and Drop-seq protocols, cells were PI stained and 120,000 live cells were sorted for each cell line by FACS to acquire an accurate equal mixture of live cells from the three cell lines. This mixture was equally split into three parts, where one part was then processed by the Chromium 10X single cell platform using the manufacturer’s (10X Genomics) protocol. The second part was processed by Dolomite Drop-seq with standard Drop-seq protocols [24]. The third part was sorted in a 384-well plate and processed using the standard CEL-seq2 protocol, with a PCR clean up ratio of 0.8 followed by 0.9. For the five cell line experiment, cells were counted using Chamber Slides and roughly 2 million cells from each cell line were mixed and processed by 10X. Three CEL-seq2 plates were sorted from the same sample, referred to as *sc*_*CEL-seq2*_*5cl*_*p1, sc*_*CEL-seq2*_*5cl*_*p2* and *sc*_*CEL-seq2*_*5cl*_*p3* in our analyses. All samples, including Drop-seq, 10X and CEL/SORT-seq, were sequenced on an Illumina Nextseq 500.

### Data preprocessing and quality control

*scPipe* was used for data preprocessing and quality control to generate a UMI-deduped gene count matrix per dataset. In general all data was aligned using *Rsubread* [26] to the GRCh38 human genome and its associated annotations, with ERCC spike-in sequences added as special chromosomes. For 10X, we processed the 5,000 most enriched cell barcodes, with `comp=3` used in the function `scPipe::detect_outliers` for quality control to remove poor quality cells. For CEL-seq2 and SORT-seq, we used the known cell barcode sequences for cell barcode demultiplexing and `comp=2` was used in the function `scPipe::detect_outliers` for quality control. For the *single cell* datasets, the population structure is informed by the cell line identity, while for the mixture data, it depends on the mixture combination. The background contamination was high for Drop-seq, so we first ran `scPipe::detect_outliers` with `comp=3` to remove outlier cells and then ran it again with `comp=2` to remove the background noise which consists of droplets that did not contain beads. The quality control metrics, including intron reads for each cell, were generated during cell barcode demultiplexing by the function `scPipe::detect_outliers`. Intron reads are defined as any reads that map to the gene body but do not overlap an annotated exon. The PCA and t-SNE results were generated using `runPCA` and `runTSNE` in the *scater* package with default parameters and perplexity was set to 30. The UMAP results were calculated using `umap` from the *umap* package.

### Data normalization and imputation

The raw counts were used as input to each normalization algorithm and all methods were blind to the biological groups. To have a fair comparison, the normalized counts from algorithms such as *BASiCS* (1.4.0) [48] and *SCnorm* (1.4.2) [2] which do not generate values on a log-scale were log2 transformed after an offset of 1 was added to the counts. The raw counts were also log2 transformed before calculating the Pearson correlation and silhouette width.

We used *edgeR* [35] (3.24.2) to calculate count-per-million (CPM) and TMM (trimmed mean of *M*-values) values. The *BASiCS* method requires spike-in genes, so we did not apply it to our datasets generated by 10X or Drop-seq which both lacked ERCC spike-ins. For *scone* [6], we set the maximum number of unwanted variation components to 0 for the removal of unwanted variation method (RUV) and ignored QC metrics. For other methods, we used their default parameters.

Data analysis using *scran* (1.8.2), *DrImpute* (1.0), *DESeq2* (1.20.0), and *SCnorm* (1.4.2) was performed with default settings. For the *RNA mixture* data, *kNN-smoothing* (2.0) was run with *k* = 16. The `size.factor` parameter was set to 1 in *SAVER* (1.1.1) to override its internal normalization procedure. *BASiCS* (1.4.0) was run with 5,000 MCMC iterations, 500 warm up iterations and a thinning parameter of 10.

Imputation methods were applied to input data generated by the different normalization algorithms, and non-normalized raw data using the `apply methods` function in *CellBench* (0.99.4).

The Pearson correlation coefficient was calculated using gene expression after normalization or imputation for samples with the same RNA mixture proportion, as these samples are replicates and any differences in gene expression should be contributed by variations in RNA amount and technical noise. We performed PCA using normalized counts and calculated the silhouette width on the first two PCs to assess whether normalization was able to preserve the known structure. For any clustering of *n* samples (here a cluster refers to a particular mixture or a cell line), the silhouette width of sample *i* is defined as

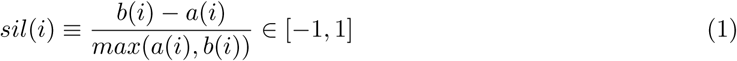

where *a*(*i*) denotes the average distance (Euclidean distance over the first two PCs of expression measures) between the *i*th sample and all other samples in the cluster to which *i* belongs to, and *b*(*i*) is calculated as below: for all other clusters *C*,

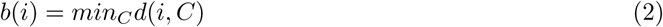

where *d*(*i, C*) denotes the average distance (the same as described above) of *i* to all observations to *C*. Methods with better performance have higher silhouette width. The function `silhouette` from the package `cluster` [31] was used to calculate the silhouette width.

### Clustering

Our comparison of clustering methods used all mixture datasets apart from cellmix5 (which is the population control). To obtain truth for the *single cell* datasets *sc*_*CEL-seq2, sc*_*10x, sc*_*Drop-seq, sc*_*10x*_*5cl, sc*_*CEL-seq2*_*5cl*_*p1, sc*_*CEL-seq2*_*5cl*_*p2* and *sc*_*CEL-seq2*_*5cl*_*p3*, we used *Demuxlet* [22], which exploits the genetic differences between the different cell lines to determine the most likely identity of each cell. The predicted cell identities in each dataset corresponded largely to clusters seen when visualising the data. Five methods, including *clusterExperiment* (2.2.0), *RaceID3* (0.1.3), *RCA* (1.0), *SC3* (1.10.0) and *Seurat* (2.3.4) were compared. Each method is used as specified by the authors in its accompanying documentation. For each dataset, the inputs for each method were normalized and imputed by different methods and the top 1,000 highly variable genes were selected using the `trendVar` and `decomposeVar` functions in *scran*. The same gene selection method was also applied in other downstream analyses such as trajectory analysis and data integration. The *Seurat* package has its own data preprocessing pipeline that takes raw UMI counts as inputs and includes normalization and gene selection (referred to as *Seurat pipe* in the results). Most methods besides *Seurat* have functions to help choose the optimal cluster numbers. Therefore two resolutions, 1.6 (*Seurat*_*1.6*) and 0.6 (*Seurat*_*0.6*) were applied to get greater or fewer clusters.

In order to compare the performance of the clustering methods, we looked at two measures: entropy of cluster accuracy, *H*_*accuracy*_, and entropy of cluster purity, *H*_*purity*_. With *M* and *N* representing the cluster assignment generated from clustering methods and annotations (ground truth), we define these measures as follows:

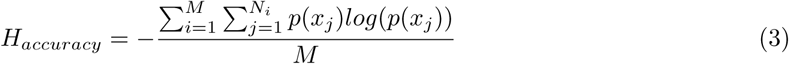

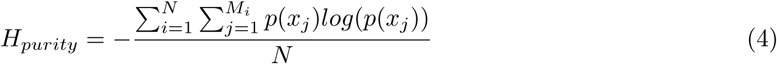

For the *H*_*accuracy*_, *M* denotes the cluster generated from a method, and *N*_*i*_ is the real clusters in *i*th generated cluster. Similarly, in the *H*_*purity*_ the *N* denotes the real clusters while *M*_*i*_ is the generated cluster for *i*th real cluster.

### Trajectory analysis

The comparison of trajectory analysis methods used all 9-cell mixture datasets (*cellmix1* to *cellmix4*) and the *RNA mixture* dataset generated by the CEL-seq2 and SORT-seq protocols. For each dataset, the gene count matrix is normalized using different normalization methods and imputation methods. The top 1,000 most highly variable genes were selected using the `trendVar` and `decomposeVar` functions in *scran*. Five methods, including *Slingshot* (1.0.0), *Monocle2* (2.8.0), *SLICER* (0.2.0), *TSCAN* (1.20.0) and *DPT* (0.6.0) were compared on the above dataset. *Slingshot* requires the dimensionally reduced matrix and cluster assignment as input. Similar to the approach described in their paper, we used PCA (`scater::runPCA`) for dimensionality reduction and Gaussian mixture model was performed on the first two PCs to obtain the cluster assignments, using `mclust::Mclust` functions. Next, the first two PCs and the clustering results were used as input for *Slingshot*. DDR-Tree, a scalable reversed graph embedding algorithm, was used for *monocle2* for dimensionality reduction and tree construction. *SLICER* applies locally linear inference to extract features and reduce dimensions. To make it easier for comparison, the samples that contains pure H2228 RNAs were selected as the root cells or root state when generating the trajectory and computing pseudotime. Then for the branching structure generated by each method, we searched for the best match to the two branches: H2228 to H1975 and H2228 to HCC827 and calculated the percentage of overlap of cells between the real path and the branch calculated by each method.

### Data integration

The main characteristics of the data integration methods applied are described in Supplementary Table 3. These analyses made use of the R packages *scran* (1.8.2) for *MNNs, Seurat* (2.3.4) for Diagonal Canonical Correlation Analysis (CCA) and *scMerge* (0.1.14). PCA and MINT analyses were performed using *mixOmics* (6.6.1) [38] and *Scanorama* (1.0) using the Python library from Hie *et al*. (2018) [16]. The input data for each analysis were the normalized and imputed results from different methods for each dataset. *scMerge* method was run in both unsupervised (referred to as *scMerge*_*us*) and supervised (*scMerge*_*s*) modes using cell identity or RNA mixtures as groups.

We calculated the silhouette width coefficient to compare the clustering performance of the different methods to combine different protocols. In the *single cell* and *RNA mixture* datasets, the clusters are defined based on either known batch/platform information or known groups. Silhouette coefficients were calculated on the first two principal components from PCA for each method that output a data matrix (*MNNs, scMerge* and *Scanorama*) or the first two resulting components for *MINT*. A high value for the batch indicates that a strong protocol effect remains, whilst a high value for the biological group information indicates that the biological variation is retained after data integration process. The kBET acceptance rate was calculated using *kBET* (0.99.5) with default parameters, with a high rate indicating homogeneous mix of samples among batches.

### Performance summary

The results from all analysis combinations including the performance scores are listed in Supplementary Table 4 and are also available from GitHub. To summarise the results of each analysis, we fitted a linear model with the performance score as dependent variable, and type of method and experimental designs as binary covariates. The coefficient of each method indicates the degree to which the method is positively or negatively associated with performance. This analysis is summarised in Supplementary Figures 4A, 4B and 8.)

Figure 6 summarises the performance across all evaluated methods. For each task, we considered a specific metric. Clustering performance was assessed with the ARI coefficient, trajectory with the correlation between pseudotime and ground truth and integration across protocols, along with a combination of normalization and imputation approaches, with the silhouette width coefficient and *kBET*. The best performance is defined as the average of the best two results for each design, and the average performance is calculated across all results. Variability refers to the variation of all results. For normalization and imputation methods, the coefficients of the linear model were scaled and shown in the heatmap to summarise performance and indicate which method yields better results in the downstream analysis compared with others.

The running time for each combination is given in Supplementary Table 5. For each method, a linear model was fitted on the running time and the number of cells on log scale. The coefficient of the number of cells indicate the scalability of the method. The scalability is classified to poor if the coefficient is larger than 2, good if the coefficient is smaller than 1 and fair if between 1 and 2. The running time for a method was regarded as poor if it was longer than 30 minutes, good if shorter than 5 minutes and fair if in between.

### Data and code availability

These data are available under GEO SuperSeries GSE118767. A summary of the individual accession numbers is given in Supplementary Table 1. The processed `SingleCellExperiment` R objects, including all code used to perform the comparative analyses and generate the figures are available from https://github.com/LuyiTian/CellBenchdata. *CellBench* is an R package developed for the benchmarking of single cell analysis methods. It contains functions and data structures that simplify the testing of combinations of analysis methods without duplicating code. In addition to a benchmarking framework, *CellBench* also provides functions to access pre-processed data objects for the samples described in this study. The *CellBench* software is available from https://github.com/Shians/CellBench.

## Acknowledgements

We thank Dr Clare Weeden and Dr Marie-Liesse Asselin-Labat for providing the cell lines used in this study. We thank Mr Jaring Schreuder and Ms Dawn Lin for assistance in conducting experiments and Mr Isaac Virshup for assistance in the data integration process.

**Funding:** This work was supported by the National Health and Medical Research Council (NHMRC) Project Grants (GNT1143163 to MER, GNT1124812 to SHN and MER, GNT1062820 to SHN), Fellowship GNT1104924 to MER, GNT1087415 to KALC, the Chan Zuckerberg Initiative DAF, an advised fund of Silicon Valley Community Foundation (grant numbers 2018-182819 to MER, 2018-182885 to KALC), a Melbourne Research Scholarship to LT, Genomics Innovation Hub, Victorian State Government Operational Infrastructure Support and Australian Government NHMRC IRIISS.

## Author contributions

LT designed, planned and performed experiments, conducted data analysis and wrote the manuscript. XD, SF, KALC, SS and AJ performed data analysis and wrote the manuscript. DAZ, TSW, AS and JSJ performed experiments. SHN and MER designed the study. MER supervised the analysis and wrote the manuscript. All authors read and approved the final manuscript.

## Competing interests

The authors declare that they have no competing interests.

---

**Supplementary Table 1.** Summary of the benchmarking datasets generated. Information on the 3 experimental designs employed, the single cell protocols used, the GEO accession numbers and parameters applied when generating cDNA libraries is listed.

| Dataset name | Experimental design | Protocol | GEO number | Protocol parameters |
| --- | --- | --- | --- | --- |
| sc_CEL-seq2 | single cells from the mixture of three cell lines | CEL-seq2 | GSM3336845 | 1X384 well plate X0.8 then X0.9 clean up for PCR products |
| sc_10x | single cells from the mixture of three cell lines | 10X Chromium | GSM3022245 | standard 10X scRNA-seq protocol |
| sc_Drop-seq | single cells from the mixture of three cell lines | Drop-seq Dolomite | GSM3336849 | standard Dolomite Drop-seq protocol |
| sc_10x_5cl | single cells from the mixture of five cell lines | 10X Chromium | GSM3618014 | standard 10X scRNA-seq protocol |
| sc_CEL-seq2_5cl_p1 | single cells from the mixture of five cell lines | CEL-seq2 | GSM3618022 | 1X384 well plate. Pooling after first strand synthesis |
| sc_CEL-seq2_5cl_p2 | single cells from the mixture of five cell lines | CEL-seq2 | GSM3618023 | 1X384 well plate. Pooling after first strand synthesis |
| sc_CEL-seq2_5cl_p3 | single cells from the mixture of five cell lines | CEL-seq2 | GSM3618024 | 1X384 well plate. Pooling after first strand synthesis |
| cellmix1 | 9 cell mixtures from three cell lines | CEL-seq2 | GSM3295024 | subsampling 1/9 from the same 384 plate. 2X0.7 clean up for PCR products |
| cellmix2 | 9 cell mixtures from three cell lines | CEL-seq2 | GSM3295025 | subsampling 1/9 from the same 384 plate. 2X0.8 clean up for PCR products |
| cellmix3 | 9 cell mixtures from three cell lines | CEL-seq2 | GSM3295026 | subsampling 1/9 from the same 384 plate. 2X0.9 clean up for PCR products |
| cellmix4 | 9 cell mixtures from three cell lines | CEL-seq2 | GSM3295027 | subsampling 1/3 from the same 384 plate. 2X0.7 clean up for PCR products |
| cellmix5 | 90 cell mixture (population controls) | CEL-seq2 | GSM3295023 | 24 samples 2X0.7 clean up for PCR products |
| RNAmix_CEL-seq2 | mixture of RNA extracted from bulk population | CEL-seq2 | GSM3305230 | 1X384 plate X0.8 then X0.9 clean up for PCR products |
| RNAmix_Sort-seq | mixture of RNA extracted from bulk population | Sort-seq | GSM3305231 | 1X384 plate X0.8 then X0.9 clean up for PCR products |

**Supplementary Table 2.**
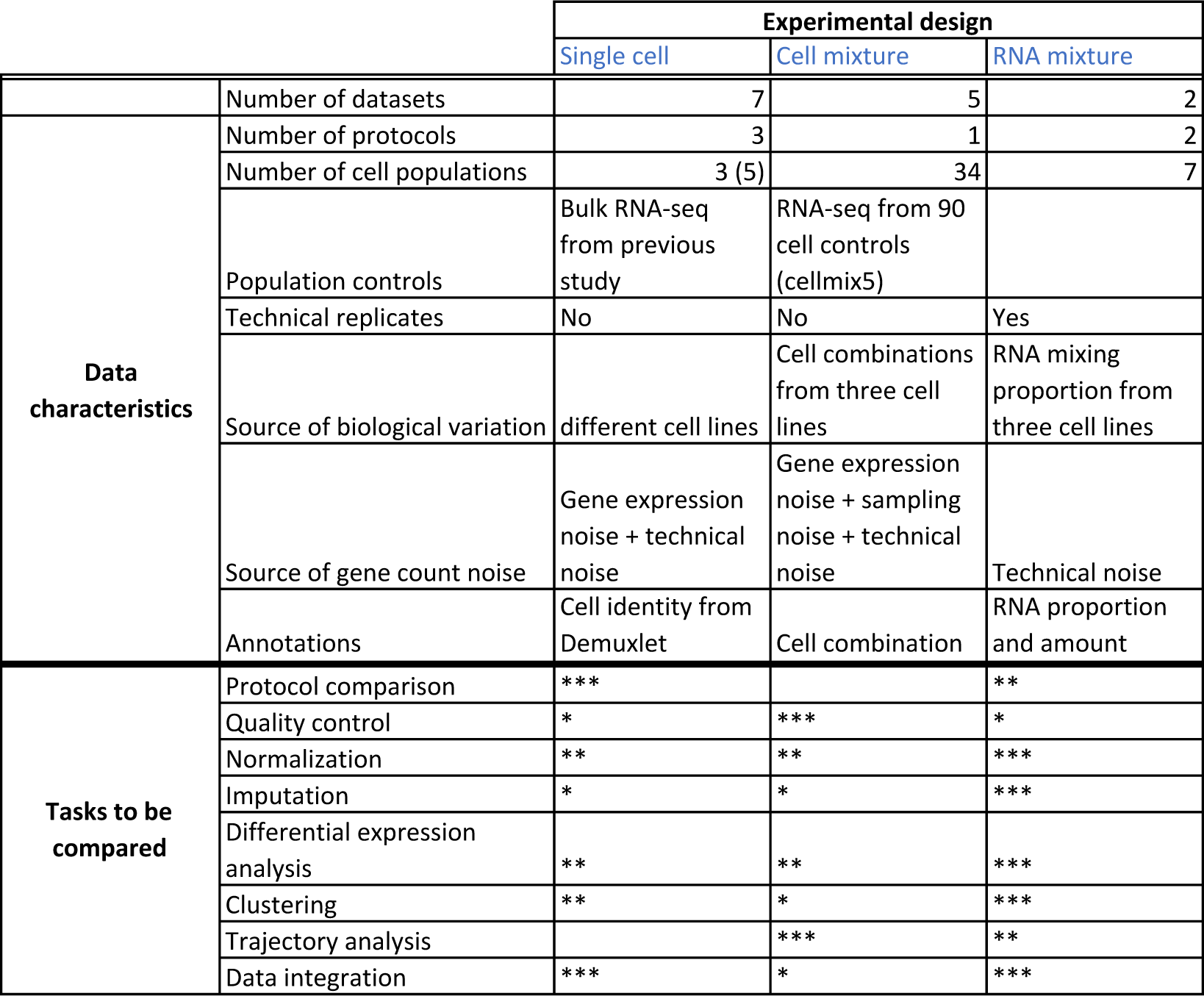
Summary of the data characteristics and data analysis tasks that can be compared by each experimental design. The suitability of each experimental design to benchmark specific tasks is indicated by the scale * < ** < *** i.e. the *RNA mixture* datasets include a dilution series which induces different dropout levels, making it an ideal dataset for comparing imputation methods.

**Supplementary Table 3.** Summary of integrative methods used to combine data from different protocols and scRNA-seq studies. Methods can be classified into batch effect correction - where a batch-corrected data matrix is output, batch effect adjustment where the batch effect is accounted for in the model and dimension reduction where components or factors summarising the batch-corrected data are output. Their hyperparameters are listed, with *italic* indicating default parameters. HVG: Hyper-Variable Genes, SEG: Stably Expressed Genes.

| Method | Correct | Adjust | Dim. reduction | # genes | Main parameters | Ref |
| --- | --- | --- | --- | --- | --- | --- |
| MNNs | ✓ |  |  | HVG | - *Number of nearest neighbors*<br>- *Bandwidth of smoothing kernel* | [13] |
| MINT supervised |  |  | ✓ | all genes | - Number of components<br>- If gene selection: Number of genes to select | [37] |
| Seurat |  | ✓ | ✓ | HVG | - Number of components<br>- Reference dataset<br>- *If multiCCA: number of iterations* | [40] |
| scMerge | ✓ |  |  | HVG<br>(+ SEG) | - Number of K-means clusters<br>- *Number of factors*<br>- *Ratio of pseudo replicates*<br>- *Distance metric* | [27] |
| Scanorama | ✓ |  |  | HVG | - *Number of HVG*<br>- *Number of nearest neighbors (NN)*<br>- *Choice of approximate kNN*<br>- *Gaussian kernel function parameter* | [16] |

**Figure S1.**
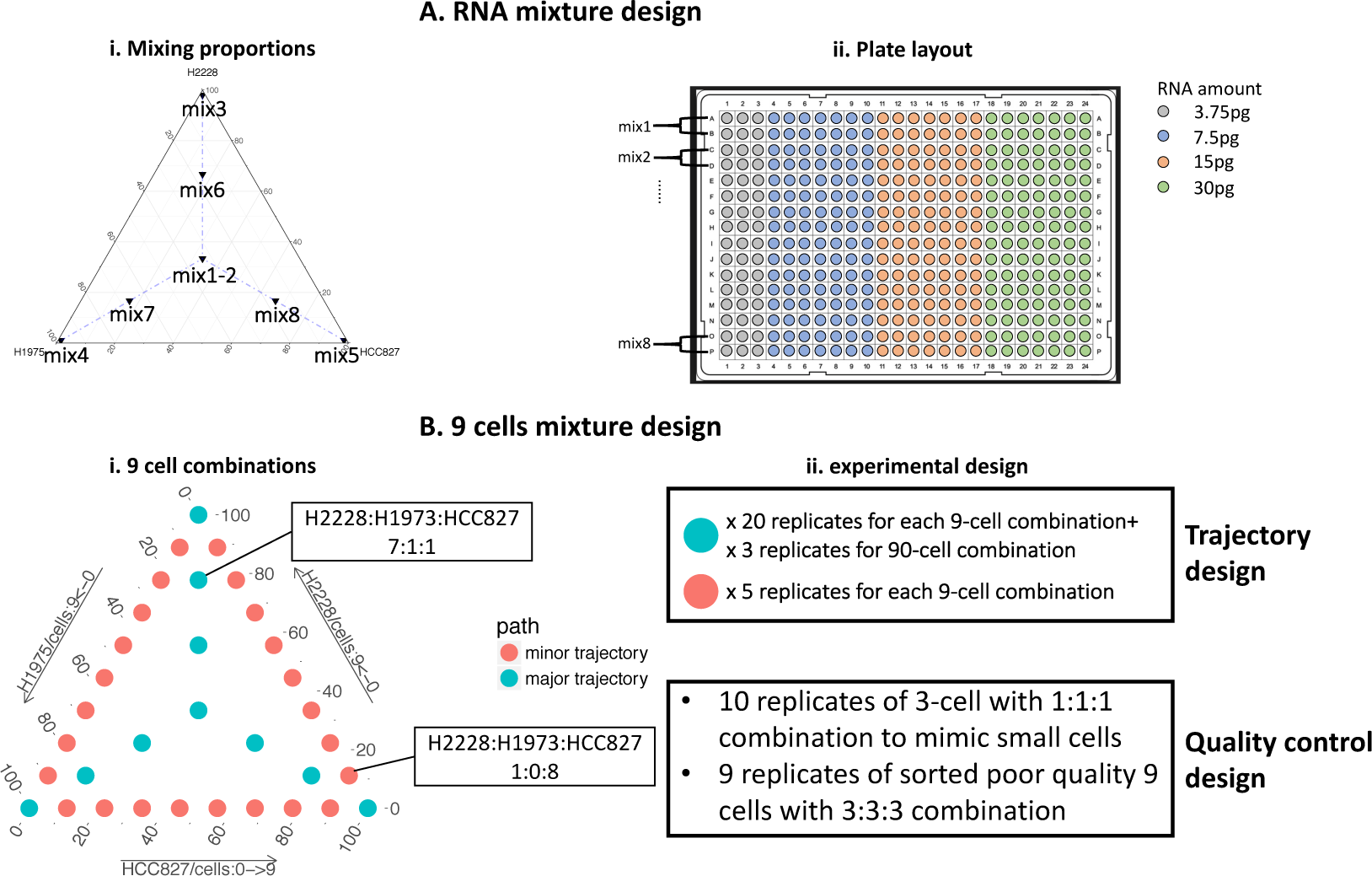
An overview of the *RNA mixture* and *cell mixture* designs from the benchmark study. (**A**) Mixing RNA extracted from bulk samples to get 8 mixtures with different proportions of RNA from 3 cell lines (**A-i**), with different amounts of RNA for each ranging from 3.75pg to 30pg (**A-ii**). (**B**) Cell mixtures, with 9 cells in total for each well in various combinations from the 3 cell lines (**B-i**). The number of replicates for each combination varies, as does the number of low quality control samples included (**B-ii**).

**Figure S2.**
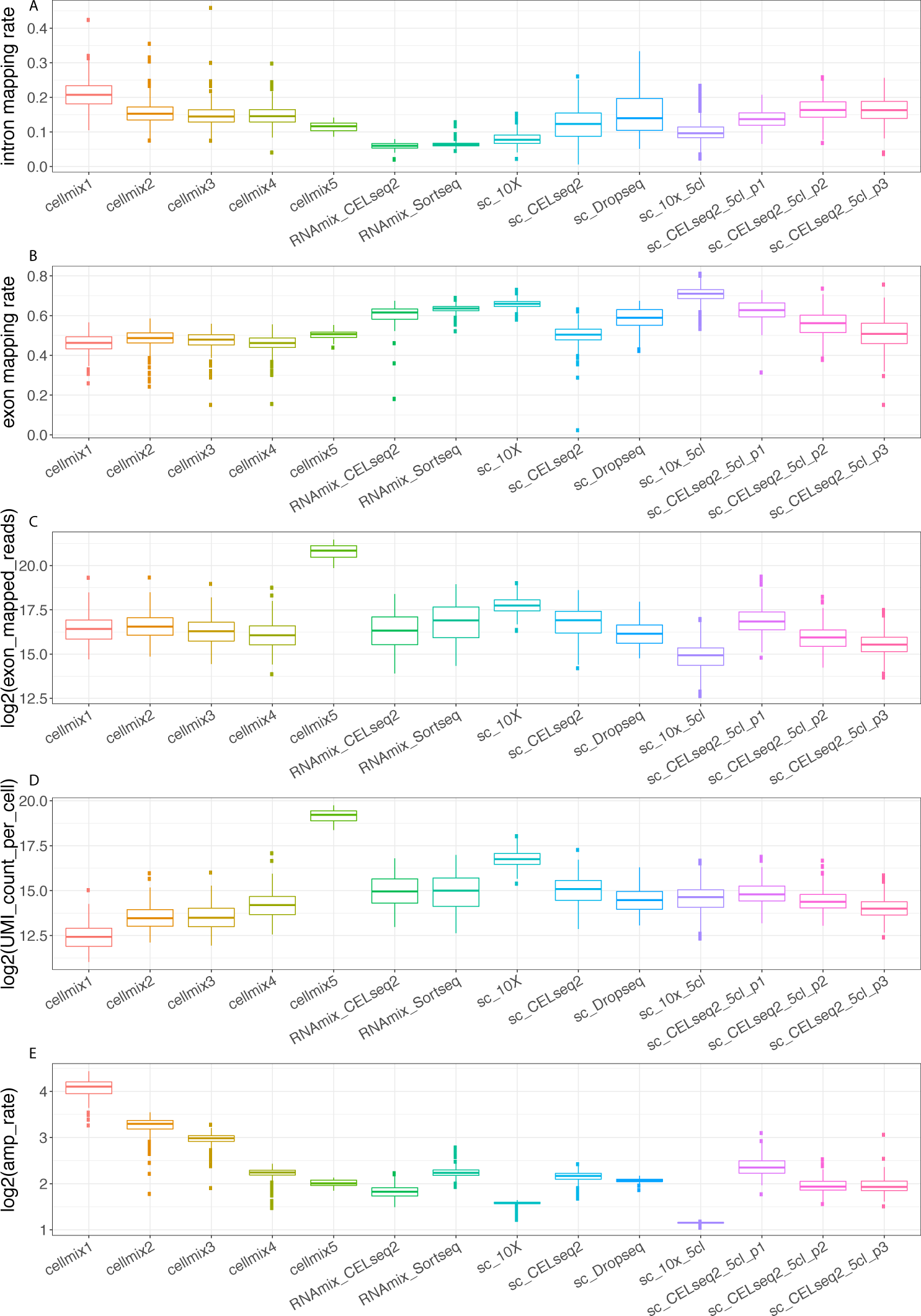
Box plots of quality control metrics for the samples from each benchmarking dataset. (**A**) The percentage of reads that map to introns. (**B**) The percentage of reads that map to exons. (**C**) The number of reads that map to exons. (**D**) The total number of counts per cell after UMI deduplication. (**E**) The amplification rate, which is defined by the ratio between the reads mapping to exons and the UMI counts after UMI deduplication. This measure reflects the library complexity.

**Figure S3.**
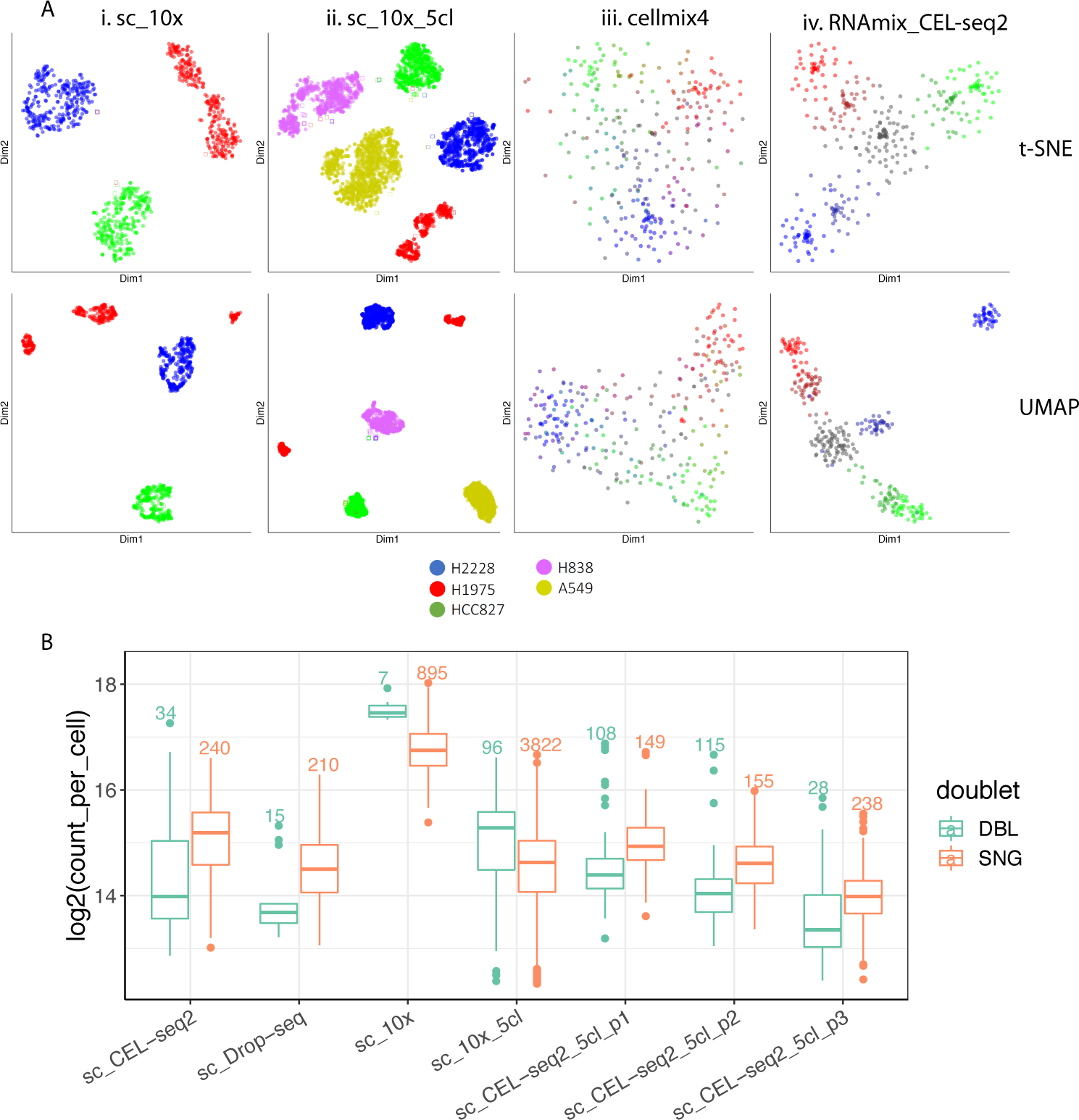
Visualisation using t-SNE and UMAP and boxplot of the number of doublets. (**A**) t-SNE and UMAP visualisation of datasets from 4 experimental designs, which are *single cell* using 3 (i) or 5 (ii) cell lines, *cell mixtures* (iii) and *RNA mixtures* (iv). Each dot represent a cell or a *‘pseudo cell’*. (**B**) Boxplot of doublets in each dataset, identified using *Demuxlet* (DBL: doublet, SNG: single cell). The number of single cells and doublets are shown on top of each box. Doublets were excluded when calculating the performance metrics.

**Figure S4.**
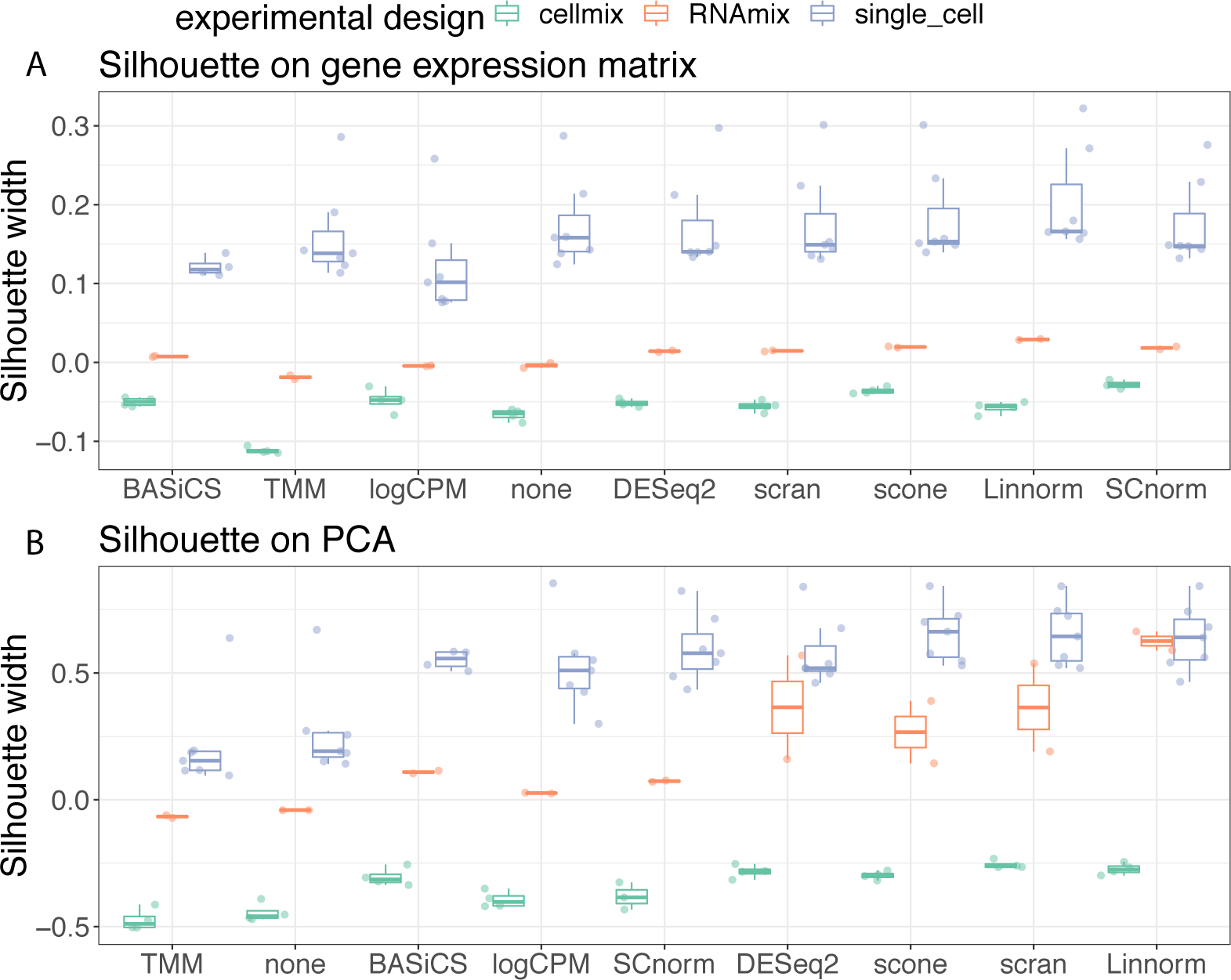
Boxplots of silhouette widths for different normalization methods. Silhouette widths calculated using the known biological groups after data have been normalized by different methods. The input to the silhouette width calculation is the distance between cells, which have been calculated using either (**A**) the gene expression matrix with 1,000 highly variable genes or (**B**) the first two PCs obtained from PCA.

**Figure S5.**
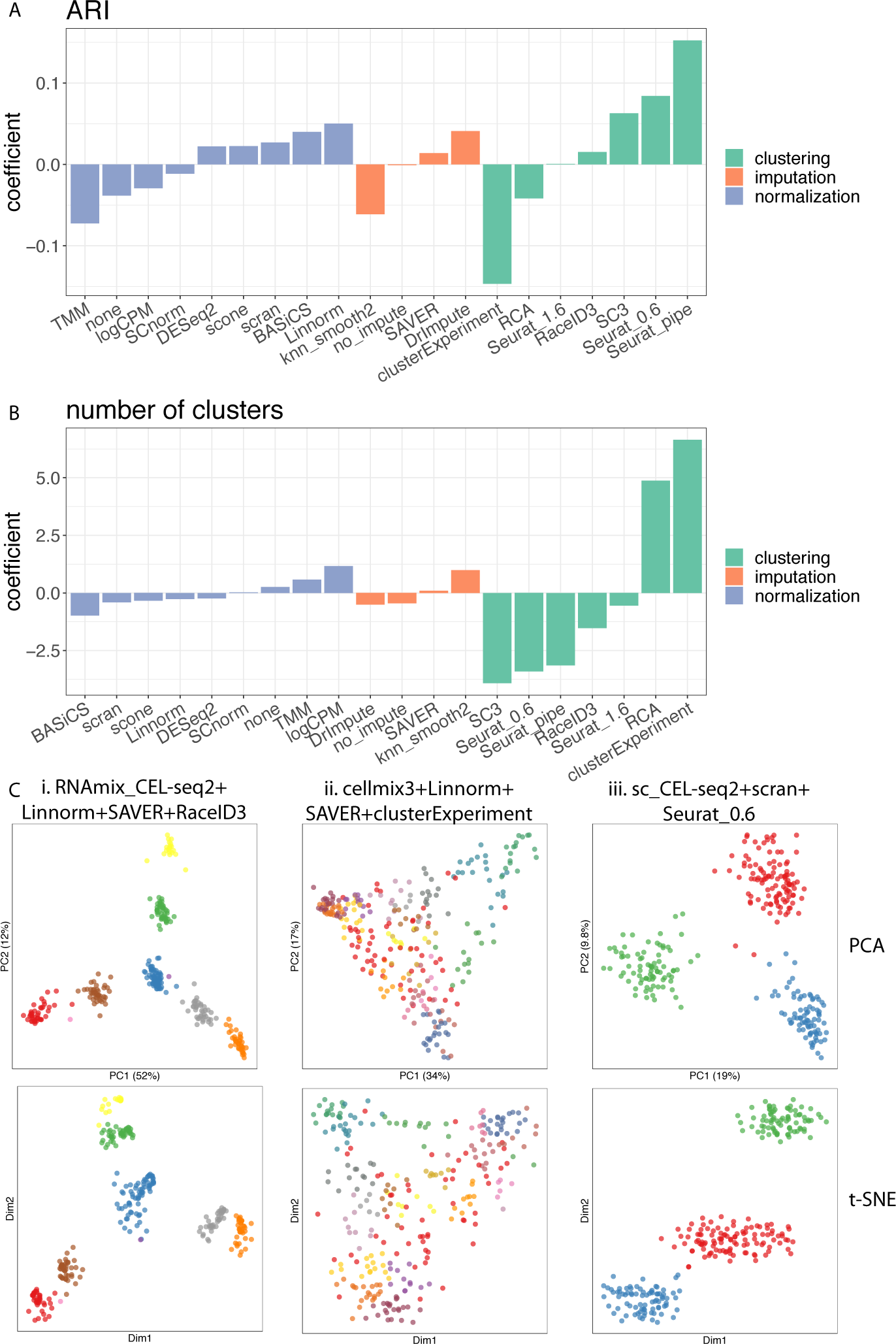
Comparisons of clustering methods using ARI and the number of clusters. (**A**,**B**) Linear models were fitted using the ARI or the number of clusters as dependent variables, and experimental design, normalization methods, imputation methods and clustering methods as covariates. The coefficients measure whether particular features have positive or negative associations with the dependent variables. (**C**) Examples of clustering results visualised by PCA (top) and t-SNE (bottom), with different colours representing the cluster assignments made by selected method.

**Figure S6.**
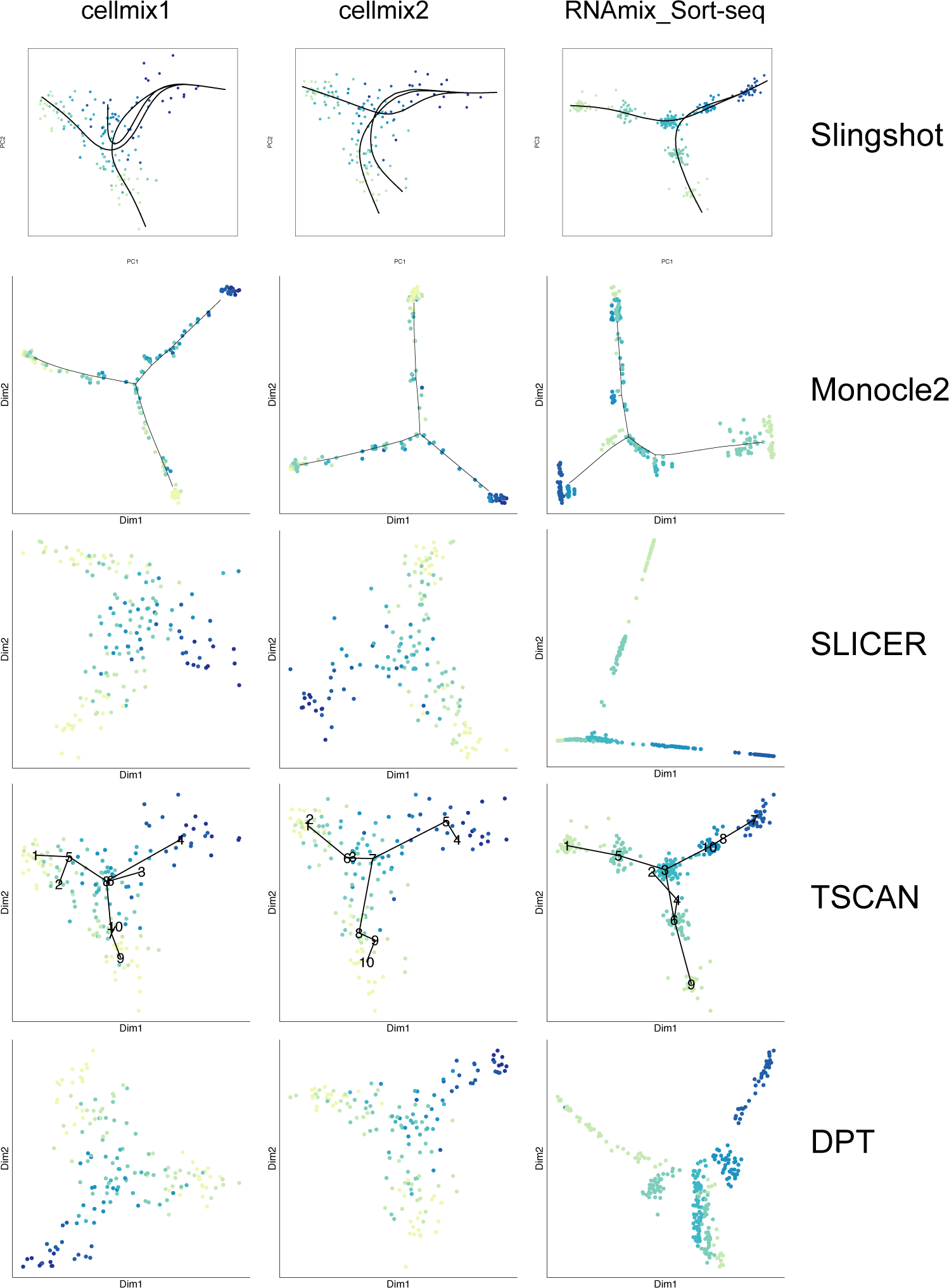
Visualisation of results from all trajectory methods evaluated in our study. Results for *cellmix1, cellmix2* and the *RNAmix_Sort-seq* analyses are shown. The dimension reduction method chosen for each method was as follows: PCA for *Slingshot* and *TSCAN*, DDR tree for *Monocle2*, diffusion map for *DPT* and LLE for *SLICER*. For this figure, data were normalized using *scran*, and the 1,000 most variable genes were input to each trajectory analysis method.

**Figure S7.**
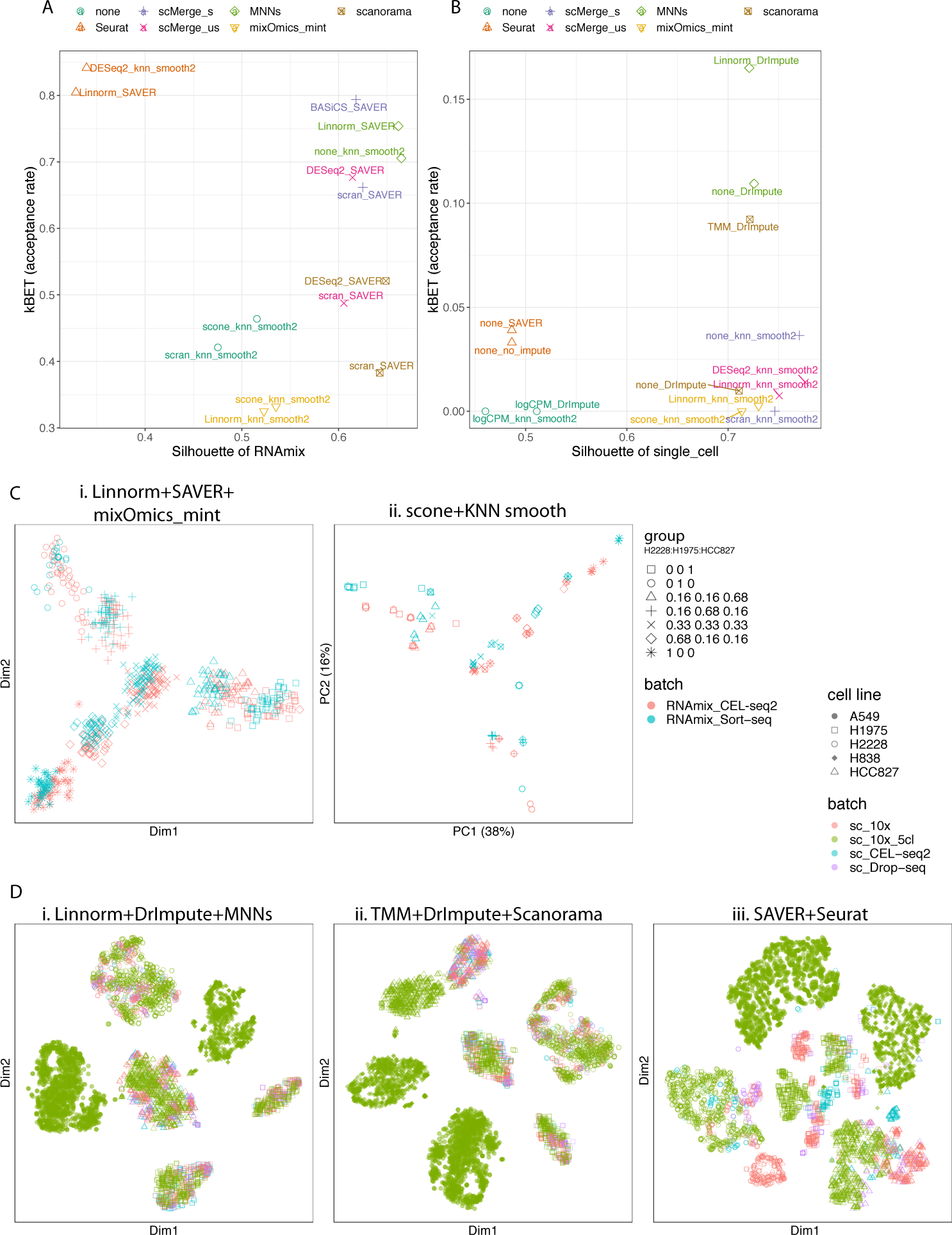
Additional data integration results for the *single cell* and *RNA mixture* datasets. (**A-B**) Sihouette width coefficient vs. *kBET* acceptance rate of each method with 2 results that have the highest sihouette width. Sihouette coefficient distance assesses the ability of a given method to group biologically similar cells together while *kBET* assesses whether different batches are homogeneous after batch effect correction. *scMerge*_*s*: supervised *scMerge*; *scMerge*_*us*: unsupervised *scMerge*) (**C**) Additional PCA and (**D**) t-SNE (perplexity = 30) visualisations where cells are coloured according to batch information (t-SNE for *MNNs* and *Scanorama* were based on batch corrected expression matrices).

**Figure S8.**
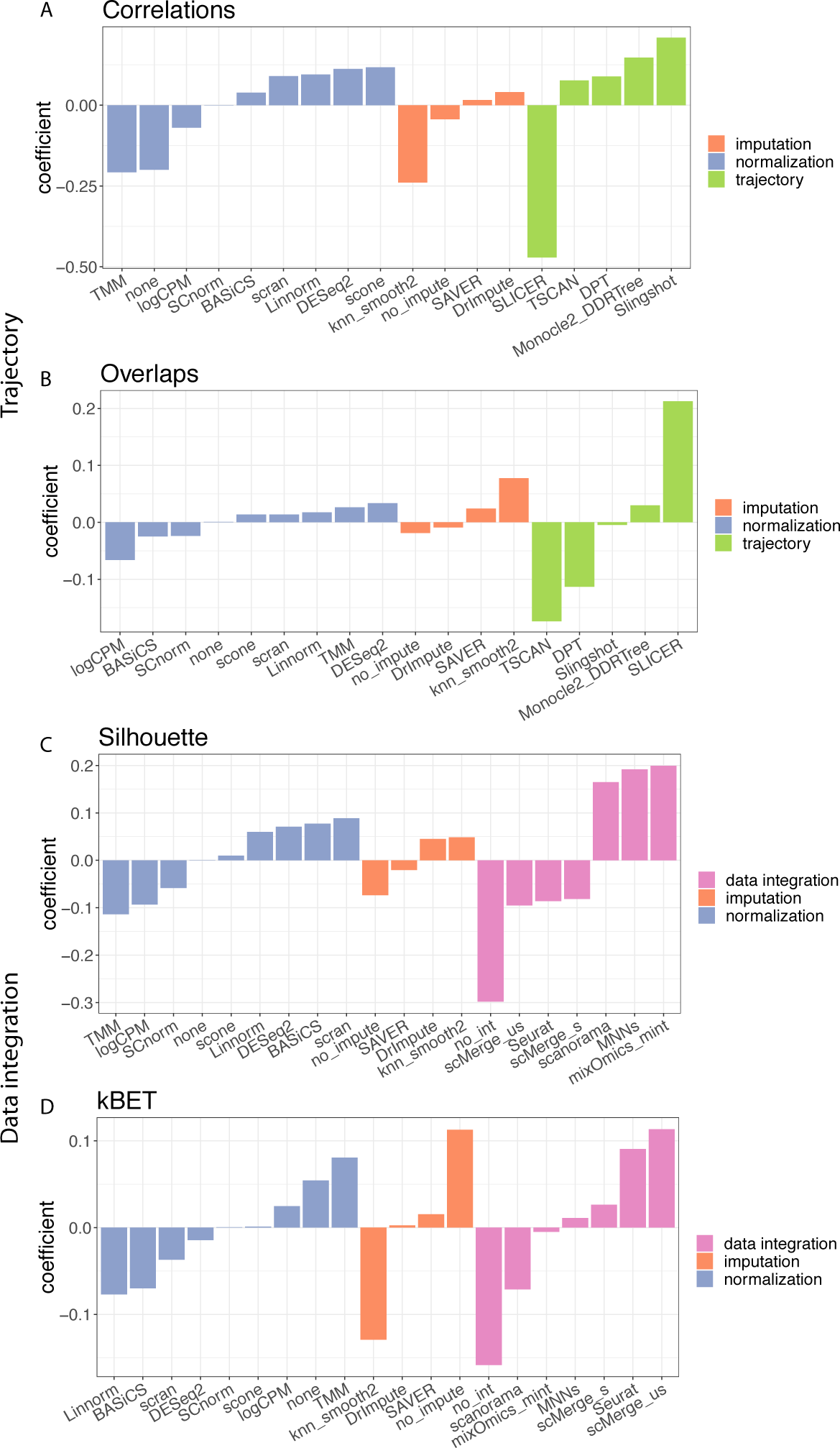
Coefficient from linear models used to quantify the impact different methods have on the trajectory and data integration results. Linear models were fitted using the evaluation metrics as dependent variables, with experimental design, normalization methods, imputation methods and trajectory analysis or data integration methods as covariates. Positive coefficients indicates that a method is positively associated with the performance metrics. The evaluation metrics used as dependent variables for each plot were: (**A**) the correlations between calculated pseudotime and ground truth; (**B**) the overlap between the calculated trajectory and the known trajectory; (**C**) the average silhouette width of the known groups and (**D**) the *kBET* acceptance rate.

